# ACORBA: Automated workflow to measure *Arabidopsis thaliana* root tip angle dynamic

**DOI:** 10.1101/2021.07.15.452462

**Authors:** Nelson BC Serre, Matyas Fendrych

**Affiliations:** Cell Growth Laboratory, Department of Experimental Plant Biology, Faculty of Sciences, Charles University, Prague, Czech Republic

**Keywords:** Root gravitropism, Image segmentation, Deep Machine Learning, Python, UNET

## Abstract

Plants respond to the surrounding environment in countless ways. One of these responses is their ability to sense and orient their root growth toward the gravity vector. Root gravitropism is studied in many laboratories as a hallmark of auxin-related phenotypes. However, manual analysis of images and microscopy data is known to be subjected to human bias. This is particularly the case for manual measurements of root bending as the selection lines to calculate the angle are set subjectively. Therefore, it is essential to develop and use automated or semi-automated image analysis to produce reproducible and unbiased data. Moreover, the increasing usage of vertical-stage microscopy in plant root biology yields gravitropic experiments with an unprecedented spatiotemporal resolution. To this day, there is no available solution to measure root bending angle over time for vertical-stage microscopy. To address these problems, we developed ACORBA (Automatic Calculation Of Root Bending Angles), a fully automated software to measure root bending angle over time from vertical-stage microscope and flatbed scanner images. Moreover, the software can be used semi-automated for camera, mobile phone or stereomicroscope images. ACORBA represents a flexible approach based on both traditional image processing and deep machine learning segmentation to measure root angle progression over time. By its automated nature, the workflow is limiting human interactions and has high reproducibility. ACORBA will support the plant biologist community by reducing time and labor and by producing quality results from various kinds of inputs.

**Significance statement:** ACORBA is implementing an automated and semi-automated workflow to quantify root bending and waving angles from images acquired with a microscope, a scanner, a stereomicroscope or a camera. It will support the plant biology community by reducing time and labor and by producing trustworthy and reproducible quantitative data.

## Introduction

Plants constantly update their internal state to adapt to the surrounding conditions and the signaling pathways involved in this process can be extremely fast. This is particularly exemplified by the root response to gravity. Root gravitropism allows to correctly orient the seedling primary root in the soil. In a matter of seconds after gravistimulation, the root columella cells sense the new gravity vector and trigger a signaling cascade to adjust the root growth towards it (1). Gravitropic response depends on rapid redirection of fluxes of the phytohormone auxin in the columella cells, on auxin flux towards to lower side of the root (2) and on auxin accumulation, perception, and response in the lower epidermis of the root (e.g., 3,4). Quantification of gravitropic performance is therefore often used as a benchmark for mutants of genes involved in auxin transport, perception, and response (2,5–7).

Technologies and methods in imaging are continuously evolving, increasing in sensitivity and spatiotemporal resolution (8–12). However, image analysis represents a bottleneck in data acquisition as it is time consuming and laborious. Turning images into numbers is particularly important to avoid cherry picking and misinterpretations. Furthermore, quantifications allow researchers to make conclusions based on proper statistical analysis. For these reasons, reproducibility of measurements is crucial; especially for tasks that are subjective (13). This is particularly the case for manual measurements of root bending as the lines to quantify the angle are usually set subjectively in software such as ImageJ. This process is then prone to human unconscious bias (14–16). Furthermore, manual measurement of root angles is a very laborious process and there is no consensus on how to measure the angle of a root bending over time. In most cases, only the angle after a certain amount of time is measured (e.g., 7,17). However, there is a growing number of scientific articles reporting gravitropism dynamics over time (e.g., 18–20). This approach is a relevant way to quantify root bending as a mutant can respond similarly to a wild type after a defined amount of time but have phenotype restricted in the first stages of gravitropic bending (21,22). In this context, implementing tools and workflows such as semi or fully automated image analysis is essential to limit the interaction of the scientists with the images and to produce trustworthy and reproducible results.

Several software tools dedicated to measuring root angles of *Arabidopsis thaliana* and other species growing on agar media are already available. However, most of them are semi-automated. Furthermore, some require specific imaging setups: PlaROM (23), RootReader 2D (24); or protocol: Kinoroot (25). Others require manual user input such as fine tuning of various parameters or manually indicating each seedlings: Rootrace (26,27). BRAT (28) is fully automated but is measuring the angle between root vector and the vertical axis of the image and thus could produce artifacts at angles bigger than 90°. These programs mostly rely on traditional image processing such as ridge detection and thresholding. However, machine learning and especially image segmentation by deep machine learning combined with the traditional methods can provide outstanding segmentation results (29–32). Deep machine learning segmentation requires prediction models trained with image libraries containing images and corresponding masks. These models can be retrained with updated libraries to expand the range of accuracy.

In parallel to scanned petri dishes, vertical stage microscopy allows high spatiotemporal observation of roots in their natural orientation in respect to gravity. Although this microscopy setup is still not a standard equipment, it continues to grow in popularity as it allows experiments such as the observation of the root gravitropic response in high spatiotemporal resolution and thus, quantification of very small bending angles (21,22). However, there is still no solution to efficiently measure root bending angle over time and measurements of small angles create noisy outputs, so it is important to have reliable measurements for small bending angles.

To provide a standard for unbiased automated measurements of root bending angles, we developed ACORBA, a fully automated workflow to measure root bending angle dynamics in a 360° space from microscope, scanner, stereomicroscope microscope or camera images. The software offers a flexible workflow based on both traditional image processing (automated or manual) and deep machine learning segmentation to measure root tip angle progression over time from images of gravitropic or waving roots. By its automated nature, the software limits human interactions and has high reproducibility. ACORBA will support the plant biologist community by reducing time and labor and by producing quality results from various types of input data.

## Results

### General presentation of the software and its workflow

ACORBA is a simple and user-friendly program written in the Python programming language (Fig. 1a). The software implements a fully automated workflow (Fig. 1b) allowing the user to measure primary root bending angle from images obtained on a microscope or flatbed scanner. ACORBA can also be used as a semi-automated approach with manual annotation of roots from various inputs such as cameras, stereomicroscopes, mobile phones.

**Figure 1:**
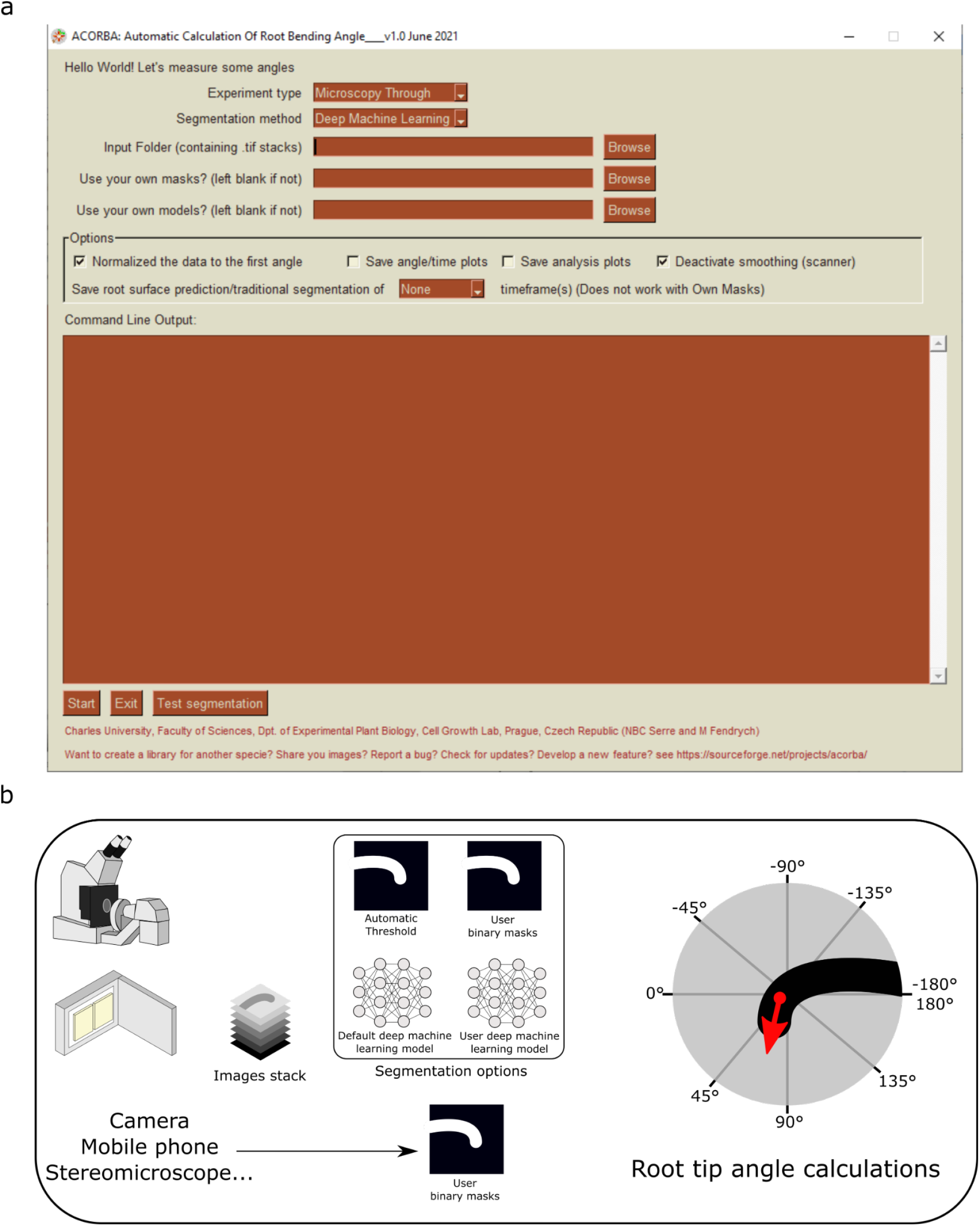
ACORBA interface and workflow. (a) ACORBA user graphic interface. (b) ACORBA workflow.

The workflow is divided into two main steps. First, the root surface is identified in the image-by-image segmentation creating a binary mask isolating the root surface(s) and root tip(s) (for microscopy images). To improve versatility, ACORBA has three possible segmentation implementations: (1) Traditional image segmentation by automatic thresholding, (2) Deep machine learning prediction models and (3) the possibility to use binary masks created by the user. The user can decide which approach is the best for a given dataset with an implemented option called “Test segmentation”. Secondly, the software automatically calculates the angle of the root tip(s). The approach used in ACORBA is the creation of a Euclidean vector which originates in the center of the root meristem and points in the direction of the actual root tip. With this method the root tip angle can be determined in a 360° space. To facilitate interpretation, the angles are exported in a +180°/- 180° space corresponding to downward/upward gravitropic bending or left/right root waving, respectively.

ACORBA includes various options including the possibility for the user to use his own deep machine learning models. These models have to follow rules described in the user manual (supplemental User Manual). This option can be used for researchers who want to analyze another species or have a large number of very specific images.

Data output-wise, the user has the option to normalize the bending angles data to the initial angle of each root. In other words, the first angle of one set of bending angles (for one root) can be automatically subtracted from the rest of the time frames to obtain relative bending angles. The raw angles are always exported as well. Users can also save the angle over time plots as well as save the analysis plots which appear during the analysis and show the vectors. These options are particularly helpful to find out the source of possible errors. Finally, to allow debugging any unusual output, the user has the possibility to save all the root predictions over time to check if the segmentation method selected is accurate for the whole stack of frames.

### Establishment of images/masks libraries and neural network architectures

Deep machine learning models are trained using libraries containing original images and their corresponding ground truth images. The lasts are binary masks (black and white pixels) in which the element to be predicted was manually annotated.

Recently, we developed a method to observe the root gravitropic response on a vertical stage microscope involving limited mechanical stress compared to the classical method of sandwiching the root between a layer of agar and the coverglass (22). In this method, which we will refer to as the “Through” method, a thin layer of ½ MS agar is casted directly on a microscopy chamber coverglass and the seedlings are placed on top (Fig. 2a). The Through method is adapted to experiments where the user is not interested in fluorescence signals in the root while the standard Sandwich method allows to have classical imaging of root fluorescence.

**Figure 2:**
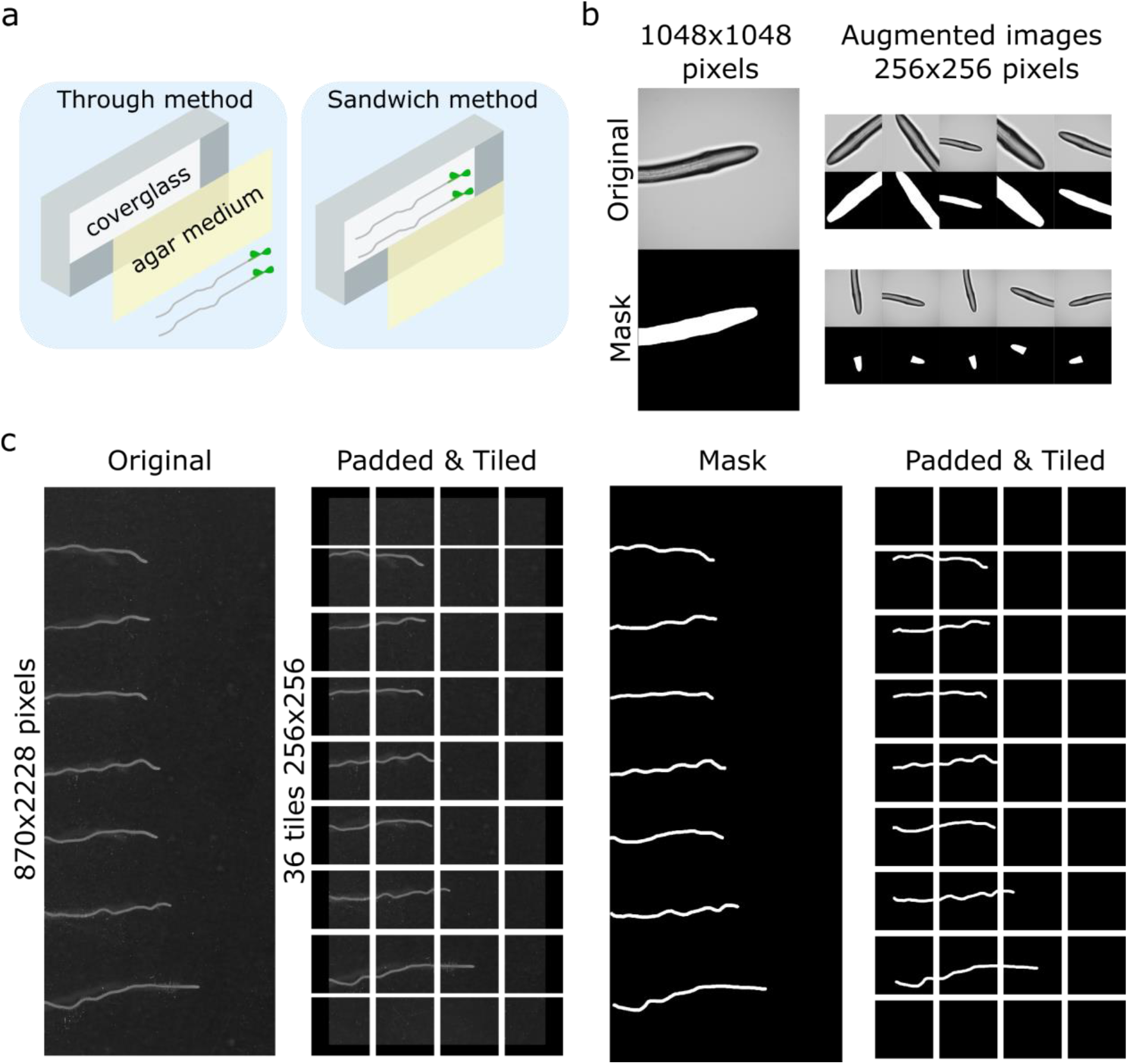
Establishment of the deep machine learning libraries. (a) Comparison of the Through and Sandwich methods. (b) Establishment of the microscopy libraries. From original images, the roots were annotated to create binary masks. Then, the original images were augmented (cropped, scaled up, rotated) and reduced to 256×256 pixels images. (c) Establishment of the flatbed scanner library. From original images, the roots were annotated to create binary masks. Then, the original images and their masks were padded with a black border to a size divisible by 256. Finally, the last images were divided into 256×256 tiles.

We created libraries and trained root surface prediction models for both microscopy methods. In parallel, we also trained root tip area detection models. Indeed, the estimation of the middle line of the root for the angle calculation requires a skeletonization step which is more accurate if restricted to a smaller area such as the root tip. Both predictions are relevant as the root surface, as a whole, is a define structure in a picture while the root tip area is a subjective zone. Prediction of the root tip area is thus, not precise enough to measure root tip angles. The original libraries created for both microscopy methods consisted of 222 and 187 images/masks for the through and the sandwich method, respectively. These numbers are in general considered small for deep machine learning training as they are unlikely to display enough diversity to obtain a prediction model with a wide range accuracy. To mitigate this problem and artificially increase the diversity and the final model accuracy, we used a method called image augmentation in which the original images and their masks are duplicated, then modified (rotation, cropping, Fig. 2b). The final libraries contained 666 and 756 images/masks for through and sandwich methods, respectively. For the root tip prediction models, the images were not cropped to not lose the overall context of the root and increase accuracy. All the models were trained with images/masks resized to 256×256 pixels for faster training and analysis. The images were obtained from several microscopy cameras (see Materials and Methods).

For images obtained with a flatbed scanner, the process was different. On one hand, skeletonization of long and thin objects is comparably easy, so we only trained a root surface prediction model. On the other hand, images of scanned square plates are large with small objects to be detected (at 1200dpi, the root thickness is approximately 10 pixels) and the images contain cotyledons and hypocotyls which are not relevant to the analysis. Therefore, cotyledons and hypocotyls were manually cropped out of the images, only leaving root parts (Fig. S2a). It was not possible to simply resize the raw images to 256×256 padded squares as this produced extreme inaccuracy in the angle measurements. To circumvent this, the raw images of various original sizes were first padded to a rectangular image that can further be divided into 256×256 tiles (Fig. 2c). The pre-library contained 340 original images/masks with 5 to 30 roots in one row obtained using commonly used Epson flatbed scanners (see Materials and Methods). The final library contained 15154 images/masks containing background or pieces of roots.

The neural networks trained in this study are modified UNET architectures (33) (Fig. S2b). This architecture is commonly used for its efficiency for semantic segmentation in biology. The network was implemented using TensorFlow implementing Keras, https://github.com/tensorflow) and the package Keras-UNET (https://github.com/karolzak/keras-unet). We used the ADAM optimizer (34) and the Jaccard index metric to quantify the prediction accuracy over the training periods. The accuracies of our trained models are presented in Table I. The root surface prediction accuracies for the microscopy methods were above 94% (For comparison with manual annotations, see Supplemental images 1-4). Given the size of the root surface, these scores are more than satisfying to allow accuracy in further angle measurements.

**Table I:** Characteristics of the trained deep machine learning prediction models.

| Model | Accuracy on library<br>(Jaccard index) | Accuracy on test images<br>(Jaccard index) | Number of<br>epochs | Model input |
| --- | --- | --- | --- | --- |
| Microscopy through<br>root surface | 94.38% | 96.43% (7 images) | 151 | 222 originals, 666*256x256<br>(augmented) |
| Microscopy through<br>root tip | 87.57% | 79.06% (7 images) | 110 | 119 originals, 666*256x256<br>(augmented) |
| Microscopy<br>sandwich<br>root surface | 96.45% | 90.35% (6 images) | 145 | 187 originals, 756*256x256<br>(augmented) |
| Microscopy<br>sandwich<br>root tip | 92.267% | 53.95% (6 images) | 71 | 187 originals, 600*256x256<br>(augmented) |
| Scanner | 72.16% | 73.03%<br>(3 images: 193 tiles) | 93 | 340 rows of seedlings divided<br>in 15154*256x256 tiles |

The accuracies for the root tips were lower, as expected. Indeed, the root tip area is not a defined area and was subjectively annotated. Overall, the trained models managed to recognize the pixel pattern created by the root tip shape. Furthermore, the root tip area detection is only intended to grossly detect the root tip in space. The scanner root surface prediction accuracy was 72%. However, consistent manual annotations of thin objects are difficult. Most of the time the final model predictions were visually more accurate than the manual annotations (For comparison with manual annotations, see Supplemental images 5-9). Deep machine learning model accuracies are also usually assessed using a test dataset, by comparing manual and prediction segmentations of images not used for the training process. The accuracies of our models on test datasets are presented in Table I and showed similar accuracies than on the libraries; with the exception of root tip predictions which were lower. These root tips were annotated separately and thus, the subjective annotations of the root tips might have been different. Regardless of the differences, the performances of the models were visually more than satisfactory (available at https://doi.org/10.5281/zenodo.5105719).

### Traditional image segmentations

As deep machine learning is unlikely to be accurate in every condition, we developed an alternative automatic solution using traditional image segmentation approaches. However, the root tip detection method is always conducted by deep machine learning as we could not find a satisfying method to detect the root tip area by traditional methods.

For the images obtained by the Through method, an automatic Otsu threshold is applied followed by a binary closing function to fill the possible holes in the binary thresholded image (Fig. 3a).

**Figure 3:**
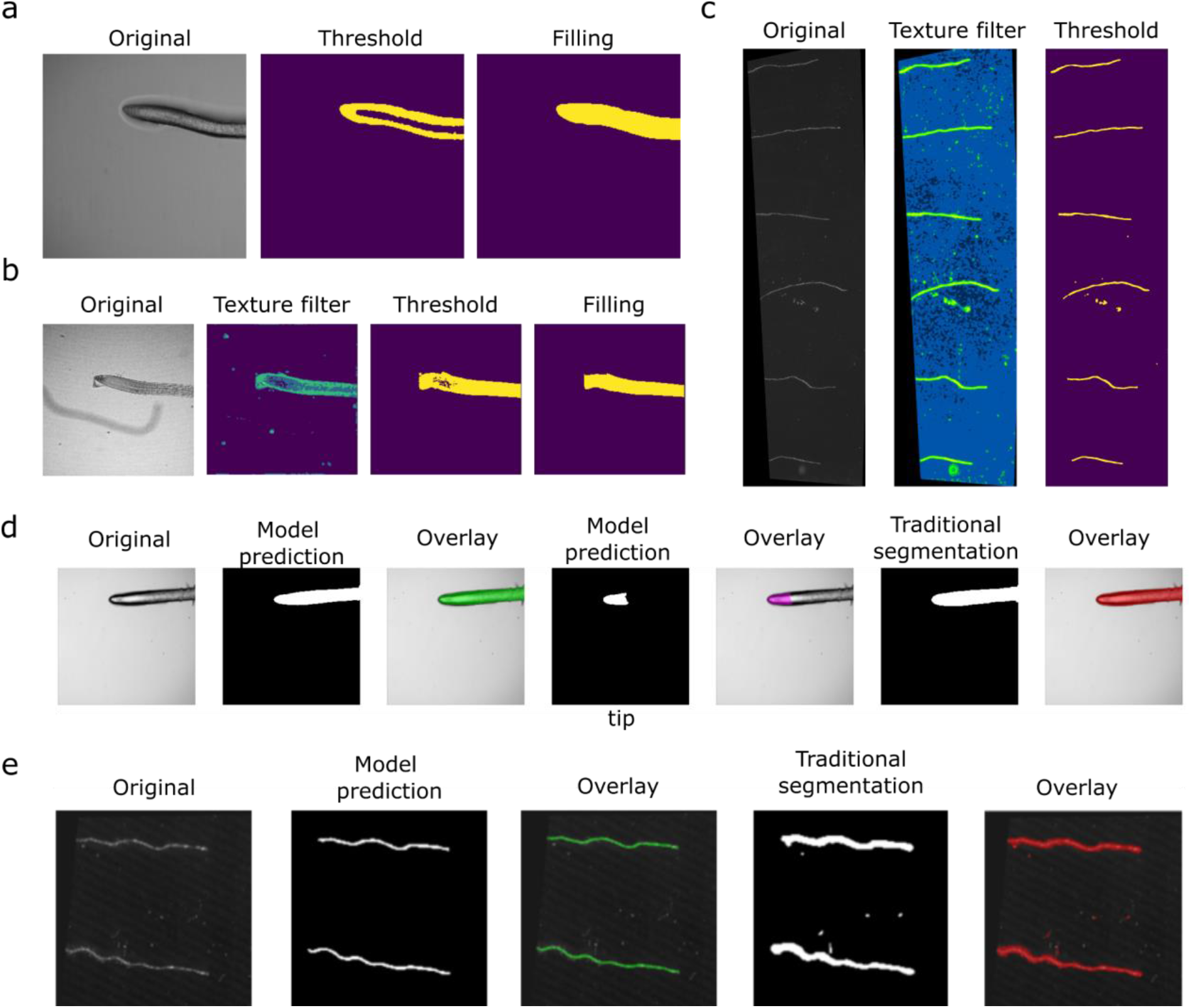
Image segmentation in ACORBA. (a) Traditional automatic segmentation of root in vertical stage microscopy using the through method. (b) Traditional automatic segmentation of root in vertical stage microscopy using the sandwich method. (c) Traditional automatic segmentation of root imaged with a flatbed scanner. (d) Example of the function Test segmentation for vertical stage microscopy using the microscopy through method. (e) Example of the function Test segmentation for scanner images.

For the Sandwich method, the image goes through a texture detection filter based on entropy (complexity of the grey levels in a given area). Then, the image is automatically thresholded and a binary closing function is applied to fill the possible holes (Fig. 3b). It is worth noting that unlike in case of the deep machine learning prediction, detached root caps and other objects with sufficient contrast are also segmented and could create angle measurements artifacts (Fig. 3b).

The method for segmentation of scanner images is similar to the ones implemented for the microscopy sandwich method (Fig. 3c).

Deep machine learning predictions and traditional methods can be compared using the “Test segmentation” option to decide which one is the most suitable for a given dataset (Fig. 3d,e). In case none of the automated methods are satisfying, the software allows the users to import their own binary masks for root surface (and root tip for microscopy). This avoids any bottleneck from image segmentation.

### Calculation of the angle for microscopy images

Even though the angle calculations are based on a root tip vector for both microscopy and scanner images, the size of the root surface and the number of roots in every image is different. Determining the root tip vector for microscopy images is a more complex process than for scanner images.

Once the segmentation of roots in microscopy images is carried out, both root surface and tip masks are preprocessed before the actual determination of the root tip vector. The first step is a series of binary pixel dilation and erosion to remove potential non-specific pixel detection conducted on both root tip and root surface masks. Then, an analysis of particles is conducted and only the largest surface (root surface/tip) is kept for analysis. Finally, to ensure that the angle orientation in space is similar for every root, the root tip is automatically oriented toward the left side of the frame.

To focus the analysis on the root tip and not the rest of the root, the root tip area is isolated. For this, a circular region of interest (40 pixels width) around the centroid of the root tip mask is created (Fig. 4a). This circular selection is used to crop and isolate the root tip area on the root surface mask. At last, this cropped area is enclosed into a circular bounding box and the root tip area perimeter is determined (Fig. 4b).

**Figure 4:**
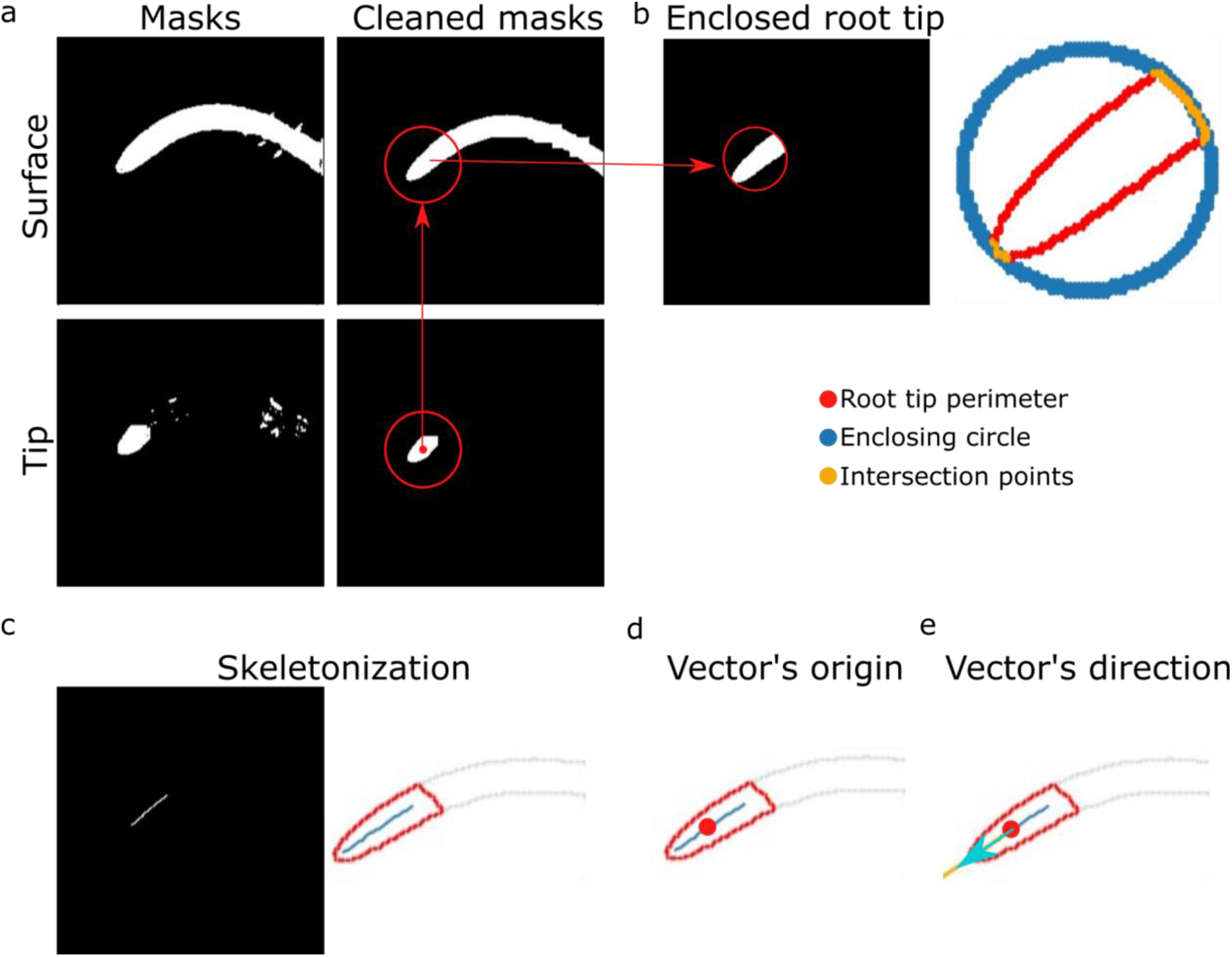
ACORBA measurements of root tip angle for vertical stage microscopy images. (a) Cleaning of binary masks and determination of the approximate position of the root tip on the root surface prediction. (b) Isolation of the root tip on the root surface prediction and determination of the intersecting points with an enclosing circle to estimate the root tip direction (c) Skeletonization of the root tip surface. (d) Determination of the middle of the skeleton corresponding to the angle vector origin. (e) Determination of the direction of the angle vector by modelling a linear regression between the estimated root tip direction (see b) and the angle vector origin. The intersection between the linear regression modelling and the root perimeter gives the direction to the vector.

The creation of the vector starts by first determining the central pixel position of the root tip area. The result of enclosing the root tip perimeter into a bounding box is the creation of two intersecting pixel clusters present both in the bounding box and the root tip area perimeter: (1) the actual root tip and (2) the root tip area shootward (Fig. 4b). These two clusters are identified by K-mean clustering. The smallest cluster central coordinates are kept for further use as it is corresponding to an estimated root tip position (Fig. 4b).

Secondly, the software goes back to the isolated root tip surface area for skeletonization (Fig. 4c). The skeleton pixels are ordered from closest to the previously determined root tip cluster to the shootward side by a closest-to-closest pixel approach. From this ordered single line of pixels, the middle pixel is set as the origin of the vector for angle calculation (Fig. 4d).

Finally, the root tip pixel (vector’s direction) precise location, is determined by creating a straight line from the middle of the skeleton to the image frame (Fig. 4e). The intersection between this line and the root perimeter is set as the root tip pixel and vector’s direction.

From this vector, the angle of the root tip is determined with the formula to determine the angle of a vector in a 360° space. However, to facilitate interpretation, 180° is automatically subtracted from the angle resulting in arbitrarily negative values for upward bending and negative values for downward bending.

This process is carried on every time frame of a stack (Supplemental video 1) and the angles are compiled into a table for export.

### Calculation of the angle for scanner images

Creation of root tip vectors from thin and long surfaces is simpler than from large surfaces. However, working with several objects in one image creates new challenges that we describe below.

Once the full binary mask is reconstructed from the 256×256 segmentation patches (Fig. 5a,b), the mask size is reduced by 40% to increase analysis speed without significantly decreasing accuracy.

**Figure 5:**
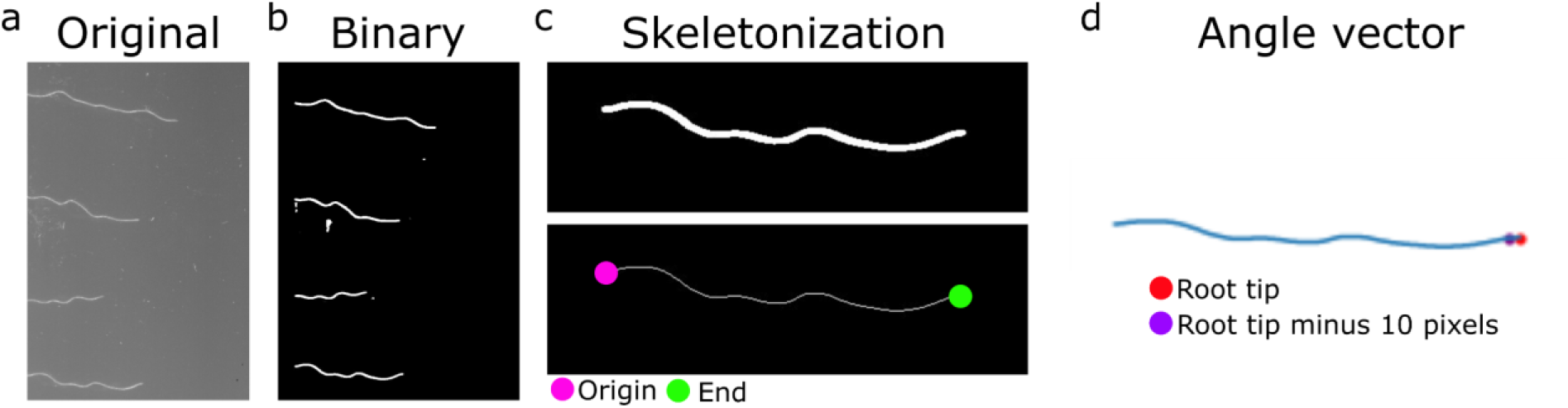
ACORBA measurements of root tip angle for scanner images. (a) Original tif stack. (b) Binary mask. (c) Individual skeletonization of every root and determination of skeleton origin and end. (d) Determination of the vector origin and direction.

The first step of the analysis is to detect every root in the images and isolate them as individuals before skeletonization which represents the roots as a single line of pixels (Fig. 5c). Next, both ends of the skeleton are detected (Fig. 5c). The identification of the shootward end and the root tip is facilitated by the mandatory step required from the users to orient the root tips towards the right side. The shootward end is determined as the leftmost coordinate and the root tip is identified as the second skeleton end. This step is followed by the reorganization of the skeleton pixels from the shootward end to the root tip by a closest-to-closest coordinate approach.

Finally, the angle vector is determined individually for each root tip. The root tip direction is determined as the root tip end identified above. The vector’s origin is set as the 10^th^ pixel on the skeleton starting from the end of the root (shootward) (Fig. 5d).

We previously stated that segmentation of several thin objects in a large image can come with challenges such as a not fully segmented root which appears as two roots. To mitigate this and ensure that all the roots are in one piece, a function calculates the distance between all the skeleton ends and origins. If one skeleton end is less than 50 pixels from another skeleton origin, the software assumes that these two pieces belong to one root and those two pieces are linked to recalculate the angles.

Similarly to the microscopy images, this whole process from reconstructing the images from tiles to the calculation of the angles is carried out on every time frame (Supplemental video 2), the angles are then compiled into a table for export.

### Evaluation of microscopy models and method

To assess the accuracy of the angle calculation method for microscopy, we first compared manual and automated analysis of the same roots bending over time in a vertical stage microscope. For the manual analysis, we used the manual equivalent of ACORBA’s method (Fig. S6). ACORBA and manual measurements, overall, produced comparable results (Fig. 6a Root 1 and 2). However, in some cases, manual measurements produced significant differences between repeated measurements of the same roots, in contrast to ACORBA which always produced the same result for one root (Fig 6a Root1 and 4). This demonstrated that ACORBA is slightly more accurate and more reproducible than the human eye and hand in manual measurements. Moreover, the analysis time by ACORBA is roughly 10 to 20 times faster (depending on the computer hardware) than the manual analysis and does not require oversight.

**Figure 6:**
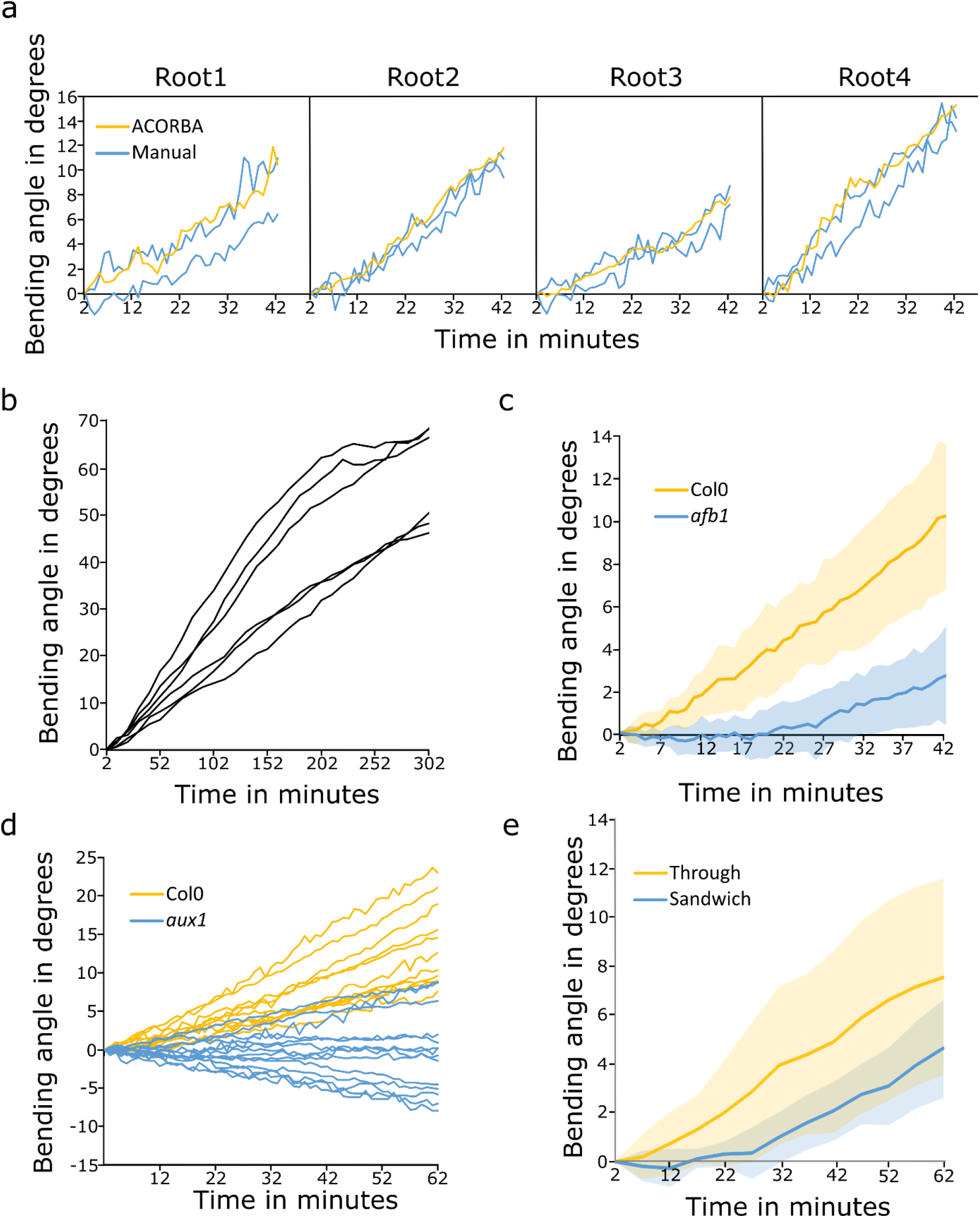
Characterization of the angle calculation method for vertical stage microscopy images. (a) Comparison of ACORBA and manual measurements of Col0 root gravitropic angles. n = 4 individual seedlings. (b) Calculation of Col0 root gravitropic angle over 302 minutes. n = 6 individual seedlings. (c) Comparison of Col0 and afb1 gravitropic bending angles. n = 9 (Col0) and 12 (afb1) individual seedlings. (d) Comparison of Col0 and aux1 gravitropic bending angles. n = 13 (Col0) and 19 (aux1) individual seedlings. Images were taken every two minutes. (e) Effect of the imaging method (through versus sandwich) on Col0 seedling gravitropic angles in a vertical-stage microscope. Represented data are mean +/- SD (shaded area). n = 10 (through) and 6 (sandwich) individual seedlings.

Next, to test whether ACORBA can measure angles bigger than the maximum 15° measured in our first experiment (Fig. 6a), we gravistimulated and measured Col0 roots bending during the course of five hours (Fig. 6b). We recorded angles up to 68°. In theory, the method would be able to measure angles higher than 180°. However, this scenario is unlikely to happen with a gravistimulated root observed with a vertical-stage microscope.

Further, we compared our recently published gravitropic data analyzed using a different method (22). In this experiment we studied the *afb1* mutant impaired in the rapid response to auxin and rapid gravitropic response. We showed that *afb1* was slower to trigger gravitropic bending with an approximative 12 minutes delay compared to the wild type Col0. The image stacks from this experiment were re-analyzed using ACORBA and showed similar results (Fig. 6c). However, ACORBA was faster than our previously published method as it is fully automated. The root tip detection method is more precise and our previous method did not allow measurements of angle higher than 90°.

Finally, to demonstrate the performances of ACORBA in another biological context, we compared the gravitropic response of Col0 and the *aux1* mutant impaired in the main auxin influx carrier in plants. This mutant is known to be agravitropic as auxin is not correctly redistributed during the gravitropic response (2). As expected, Col0 initiated bending toward the gravity vector with recorded downward angles ranging from 5 to 21° (Fig. 6d). In contrast, *aux1* roots displayed a typical agravitropic behavior with growth orientation independent of the gravity vector with roots bending upward, downward or not bending, again well analyzed by ACORBA (Fig. 6d).

We previously described two methods used in our laboratory to assess the gravitropic response using a vertical-stage microscope (Fig. 2a). Here, we used ACORBA to compare the dynamics of root bending imaged with the Through and Sandwich methods. This experiment showed that the roots imaged with the through method are, overall, bending faster than the roots imaged with the sandwich method (Fig. 6e). This confirmed that the through method is allowing a better gravitropic response probably by providing less mechanical stress and unobstructed growth.

This set of experiments allowed us to confirm that ACORBA can be used to accurately determine the bending angles of roots imaged with a vertical-stage microscope.

### Evaluation of the scanner model and method

We progressed further with the evaluation of the scanner angle calculation. We first compared ACORBA and manual measurements (method described in Fig. S6a) of root bending on agar plates measured over 15 hours after gravistimulation (Fig. 7a). We showed that ACORBA and manual measurements were strikingly similar. However, ACORBA produced more noisy data. Nevertheless, in most cases the bending angles of roots in plates are averaged and the overall average and error bars between methods were almost identical. Furthermore, the analysis time is considerably shorter with ACORBA (up to 10 times depending on hardware) and does not require oversight.

**Figure 7:**
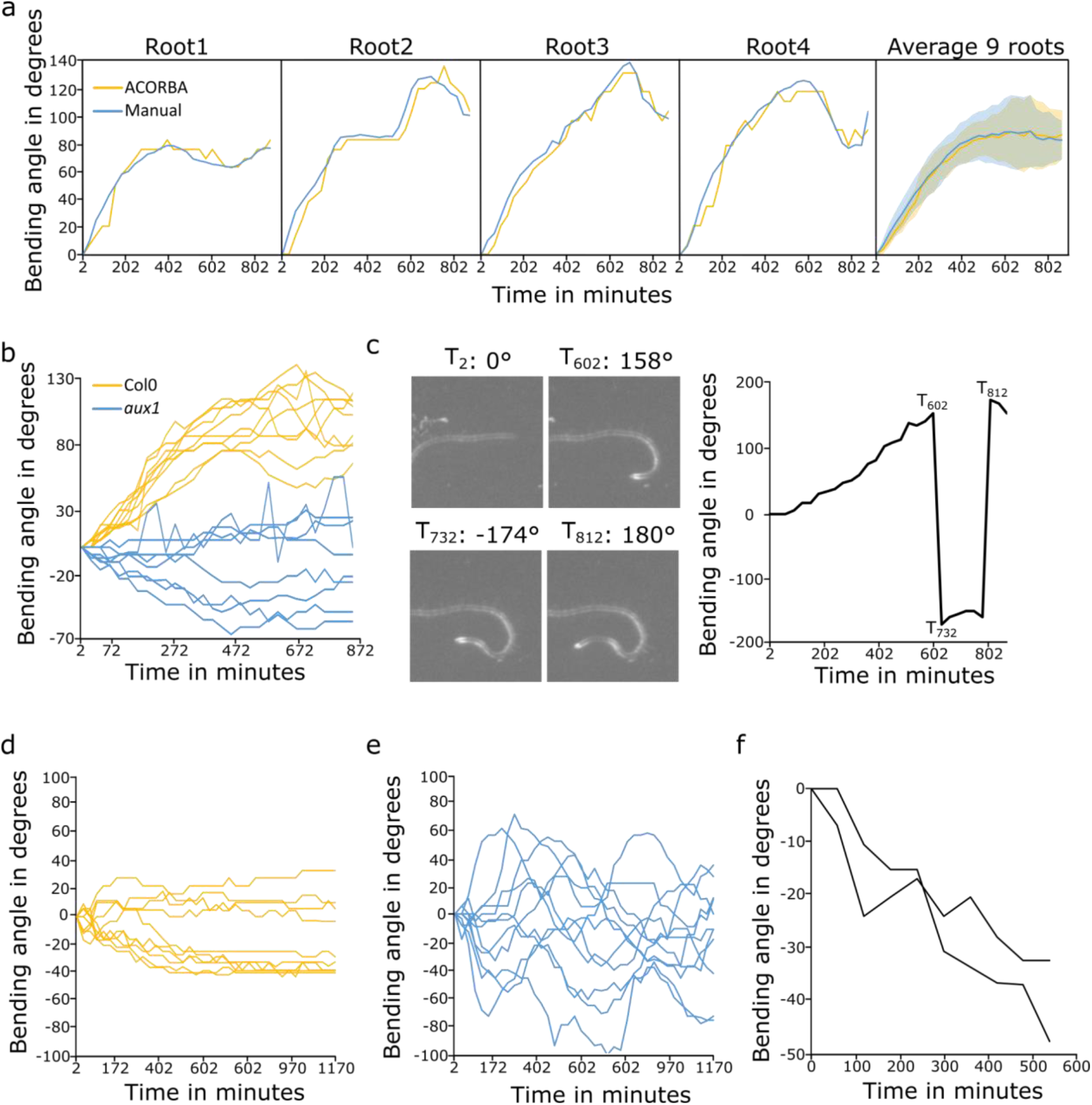
Characterization of the angle calculation method for flatbed scanner images. (a) Comparison of Manual and ACORBA measurements of Col0 gravitropic bending angles. n = 9 individual seedlings. (b) Comparison of Col0 and aux1 gravitropic bending angles. n = 9 (Col0) and 8 (aux1) individual seedlings. (c) Measurement of root tip angles during horizontal root curling. (d) Effect of no sucrose and (e) 3% sucrose on Col0 vertical root waving. n = 10 (0% sucrose) and 11 (3% sucrose) individual seedlings. (f) ACORBA semi-automated measure with manual binary masks of root tip angles of Col0 seedling growing horizontally and observed under a stereomicroscope. Images were taken every thirty minutes.

Next, we also compared Col0 and the agravitropic mutant *aux1* responses and showed that while Col0 bent downward up to 130° over 15 hours, the mutant bent slightly downward or upward, showing a typical agravitropic behavior (Fig. 7b).

To push the limits of the software, we imaged a root curling, a phenomenon appearing when agar plates are horizontally positioned. In this scenario, the deep machine learning segmentation showed untrustworthy accuracy above 158° bending. However, the automatic traditional segmentation method and the angle calculation method showed that ACORBA can measure relative angles up to 180°. Passing the 180° threshold up/downward bending resulted in a complex output, with difficult interpretation as the root tip is changing horizontal direction (Fig. 7c). The complex interpretation is a direct consequence of working in a +180/-180° space instead of a 360° one. However, this situation is not common and the gain in interpretation facility obtained by working in a +180/-180° space for regular gravitropic experiments overcomes this specific limitation.

To demonstrate the software ability to measure angle of waving roots, we imaged seedlings growing vertically in 1/2MS without or with 3% sucrose, known to induce waving (For review, 34) for 20 hours. The seedlings transferred from 1% sucrose to no sucrose showed almost straight root growth after 6 hours (Fig. 7d). On the other hand, seedlings transferred to 3% sucrose medium displayed heavy waving with a waving amplitude of approximately +/-60° (Fig. 7e).

Finally, to illustrate the semi-automated use of the software, we imaged seedlings growing in agar plates using a stereomicroscope and a mobile phone (16 megapixels resolution) (Fig. 7f). After resizing the images, so that the roots were approximately 10 pixels in width (similar to 1200 dpi scanned images), we prepared the binary masks manually in ImageJ (Method in Supplemental User manual, Fig. S7a,c). It is worth noting that seedlings from the mobile phone image and the stereomicroscope were also correctly segmented by the deep machine learning model Fig. S7b,d). In these conditions ACORBA was able to measure single time frame angle from mobile phone image (Table II) and angle dynamic of horizontally growing seedlings observed with a stereomicroscope (Fig. 7f).

**Table II:**
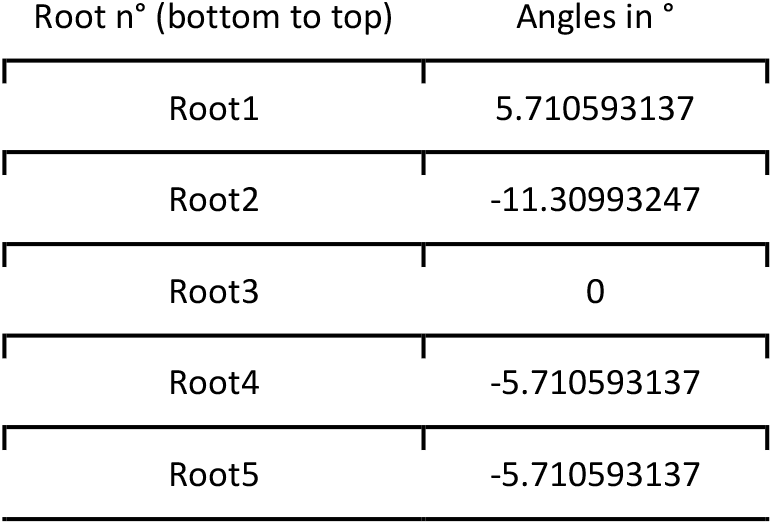
Single frame root angle from mobile phone image.

## Discussion

ACORBA is, to our knowledge, the only software dedicated to automated or semi-automated measurements of root bending angles dynamics from various image inputs. We demonstrated that it can quantify root bending towards gravity, root waving in vertical conditions or root curling in horizontal condition. The software implements semantic image segmentation with deep machine learning and can be fully automated which is still a rare case in plant biology image analysis. We implemented an innovative, deep machine learning based method to detect the root tip area in microscopy images even if the root is over-bending or spiraling over itself. This method is more accurate than our previously published method (22) and is adapted to angles higher than 90°.

The models provided in the software were trained on libraries containing images taken from different inputs to fit other laboratory setups. Furthermore, the models can still be retrained with contributions from the scientific community to expand the range of accuracy. On the other hand, as long as the input format follows our model input pre-formatting, custom deep machine learning prediction models can also be implemented for various applications. Otherwise, to avoid analysis bottlenecks and flexibility at the root segmentation step, the software can also do traditional automatic segmentation or can be semi-automated by using manually annotated roots. This opens the range of usage to researchers working with different species than *Arabidopsis thaliana*, specific image types (e.g., images from a camera, mobile phone or stereomicroscope).

The software showed similar or better accuracy than the manual measurements of root angles from microscope. For the scanner images, we showed that ACORBA produced strikingly similar measures as the one obtained by manual measurements. Moreover, ACORBA measurements are highly replicable in contrast to the manual subjective approach.

We demonstrated that it can reproduce and quantify already published data such as the delayed gravitropism of *afb1* (22), *aux1* agravitropism (2) and sucrose-induced waving in vertical conditions (35). We showed that the software was producing hard to interpret data passing the 180° angles which was induced by curling in horizontal conditions. These conditions are unlikely to happen in regular vertical bending and this experiment had for only purpose to demonstrate the limitations of interpretation passed this threshold.

ACORBA was not designed to measure angles from a single timeframe. However, by duplicating the single frame to obtain a two frames stack prior to analysis, we showed that it can be used for single time frame angle measurements.

A direct benefit but also disadvantage of the fully automated approach is that the user has less control on the measurement. This is especially striking when a root or row of seedlings displays unforeseen characteristics. Nevertheless, with the export related to the analysis steps (original/prediction overlay and analysis graphics), users can quickly identify problems and, most of the time, fix it by image preprocessing (e.g., remove an agar bubble or a detached root cap, see user manual for troubleshooting).

The program was written in the Python programming language which is not as fast a compiled language like C++ but offers very good support and modules for machine learning and image processing.

In this context, ACORBA offers a broad range of measurements possibilities by producing unbiased and highly replicable datasets by automatization of a time and laborious process.

## Experimental procedures

### Plant material and growth conditions

Wild-type *Arabidopsis thaliana* ecotype Columbia (Col0) and *aux1* (SALK_020355) were used in this study. The genotype of *aux1* was verified by PCR-genotyping using the following primers. SALK LB1.3 primer (ATTTTGCCGATTTCGGAAC), aux1-R (AGCTGCGCATCTAACCAAGT) (and the aux1-L primer). Seeds were surface sterilized by chlorine gas for 2 hours (36). Seeds were sown on 1% (w/v) agar (Duchefa) with ½ Murashige and Skoog (MS, Duchefa, 1 % (w/v) sucrose, adjusted to pH 5.8 with KOH 1M, and stratified for 2 days at 4°C. Seedlings were grown vertically for 5 days in a growth chamber with 23°C by day (16h), 18°C by night (8h), 60% humidity, light intensity of 120 μmol photons m-2 s-1.

### Plant imaging

In microscopy Sandwich method, seedlings were placed onto a thin layer of ½ MS medium placed inside a custom 3D printed chambered coverglass (24 x 50mm). The seedlings were allowed to recover vertically for at least 30 minutes before gravistimulation. In the Through setup, the roots were growing unobstructed on the surface of the agar and the imaging was performed through the coverglass and the agar.

Imaging was performed using a vertical stage (8) Zeiss Axio Observer 7 coupled to a Yokogawa CSU-W1-T2 spinning disk unit with 50 μm pinholes and equipped with a VS-HOM1000 excitation light homogenizer (Visitron Systems). Images were acquired using the VisiView software (Visitron Systems) and Zen Blue (Zeiss). We used the Zeiss Plan-Apochromat 20x/0.8 and Plan-Apochromat 10x/0.45 objectives. Brightfield signal was retrieved using a PRIME-95B Back-Illuminated sCMOS Camera (1200 x 1200 px; Photometrics), Orca Flash 4.0 V3 (2048 x 2048 px; Hamamatsu) and the camera from a LSM 700 confocal microscope (Zeiss).

For the scanner, seedlings were transferred to ½ MS, 1% sucrose (unless specified otherwise) and imaged every 30 minutes using an Epson Perfection v370, v600 or v700 flatbed scanner. The procedure followed to automate the scanning process is described in the supplemental User Manual.

### Programming

All the scripts used in this study were written in the Python programming language. The latest versions of the ACORBA software and user manual can be found at: https://sourceforge.net/projects/acorba. The software is maintained by NBCS.

The models were trained using Google Colaboratory Pro GPU accelerated servers. The software was programmed and analysis were conducted on a Windows 10 computer with an Intel^®^ Core™ i5-7300HQ 2.50GHz processor, a CUDA enabled NVIDIA GeForce GTX 1060 Max-Q design 6Go GPU and 16Go of RAM.

This work was partially inspired by Sreenivas Bhattiprolu Python image analysis and machine learning tutorials (https://github.com/bnsreenu/python_for_microscopists).

The Python dependencies used to create and run ACORBA are listed below: Scikit-Image (0.18.1), Scikit-Learn (0.24.1), TensorFlow (2.4.1), Keras (2.4.3), Keras-unet (0.1.2), OpenCV (4.5.1.48), PySimpleGUI (4.34.0), Matplotlib (3.3.4), Pandas (1.2.3), Numpy (1.19.5), Fil Finder (1.7) and Astropy (4.2.1). These dependencies are all common modules which can be installed with the Python pip command.

The Python scripts were transformed as .exe files with the module Auto PY to EXE (2.8.0) to make the scripts portable for Windows x64.

The whole software was compiled as a setup .exe file for Windows using the software Inno Setup Compiler (6.1.2).

## Supporting information

Supplemental User Manual

Supplemental image 1

Supplemental image 2

Supplemental image 3

Supplemental image 4

Supplemental image 5

Supplemental image 6

Supplemental image 7

Supplemental image 8

Supplemental image 9

Supplemental image 10

## Acknowledgements

The authors would like to thank Shiv Mani Dubey, Monika Kubalová and Denisa Oulehová for their image contributions to establish the prediction model libraries and their valuable inputs and software beta testing. Eva Medvecká for technical support. Sreenivas Bhattiprolu for his free and online Python image analysis and machine learning tutorials.

## Availability of data and materials

The latest versions of ACORBA software training annotated libraries, source code, examples, image pre-processing scripts, deep machine learning model training Jupyter notebooks and user manual maintained by NBCS are available at https://sourceforge.net/projects/acorba/. The raw microscopy and scanner stacks used in this paper are available at ZENODO (https://doi.org/10.5281/zenodo.5105719). The analyzed results are supplemented (Supplemental data).

## Author contributions

NBCS and MF conceived the project and wrote the manuscript. NBCS programmed the software, measured and analyzed the data.

