## Supplemental User Manual for "ACORBA: Automated workflow to measure *Arabidopsis thaliana* root tip angle dynamic"

### ACORBA:

### Automatic Calculation Of Root Bending Angle

Hello World! Let's measure some angles

#### USER MANUAL v1

**NBCS, June 2021**

### 1. General information

#### a. What is ACORBA?

ACORBA, standing for Automatic Calculation Of Root Bending Angle, is a software allowing you to measure root angles overtime from microscopy and scanner images. It was originally developed to measure *Arabidopsis thaliana* root bending angle during gravitropic experiment but can, with some tweaks, probably work with different species. This software allows you to use different methods for segmenting your roots:

- Deep Machine learning with default prediction models
- Deep machine learning with custom prediction models (models can be created by the user or in collaboration with us, see Custom models)
- Automatic traditional segmentation (Background cleaning and Threshold method)
- Your own segmentation images (see the page see Own segmentations)

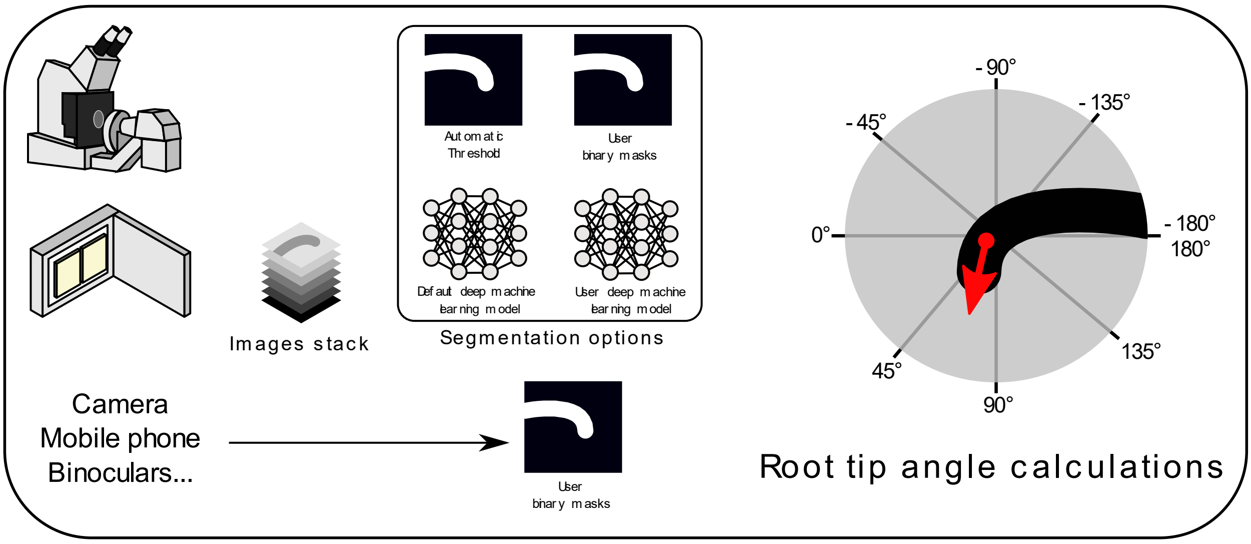

ACORBA workflow

ACORBA is fully developed in the Python language (v3.8, python.org) and compiled to an executable with the PyInstaller module (https://pyinstaller.readthedocs.io/).
The project is hosted on SOURCEFORGE and is implementing TensorFlow with Keras for deep machine learning image segmentation.

https://sourceforge.net/projects/acorba/

#### b. Contact

For troubleshooting, model or feature implementation requests, participation in development or any others information, please open a forum thread or a ticket at https://sourceforge.net/projects/acorba/

#### c. License

ACORBA software, methods and python scripts are protected under the Creative Commons Attribution-NonCommercial 2.0 Generic (CC BY-NC 2.0) license.

This is a human-readable summary of (and not a substitute for) of https://creativecommons.org/licenses/by-nc/2.0/legalcode

*Disclaimer.*
*You are free to:*
*Share — copy and redistribute the material in any medium or format*
*Adapt — remix, transform, and build upon the material*
*The licensor cannot revoke these freedoms as long as you follow the license terms.*

*Under the following terms:*
*Attribution — You must give appropriate credit, provide a link to the license, and indicate if changes were made. You may do so in any reasonable manner, but not in any way that suggests the licensor endorses you or your use.*

*NonCommercial — You may not use the material for commercial purposes.*

*No additional restrictions — You may not apply legal terms or technological measures that legally restrict others from doing anything the license permits.*

*Notices:*
*You do not have to comply with the license for elements of the material in the public domain or where your use is permitted by an applicable exception or limitation.*
*No warranties are given. The license may not give you all of the permissions necessary for your intended use. For example, other rights such as publicity, privacy, or moral rights may limit how you use the material.*

*For more details, see https://creativecommons.org/licenses/by-nc/2.0/legalcode*

### 2. Requirements and Installation

#### a. Requirements

- Microsoft Windows x64 bits version (only tested on Windows 10).

- ACORBA requires an installed version of Microsoft Visual C++ redistributable for Windows x64 bits. See: https://support.microsoft.com/en-us/topic/the-latest-supported-visual-c-downloads-2647da03-1eea-4433-9aff-95f26a218cc0

- Minimum hardware:
  - 8Go RAM (! avoid handling large stacks !)
  - 4 cores CPU with base clock >2.90GHz

- Recommended hardware:
  - >=16Go RAM
  - >6 cores CPU with a base clock >2.90GHz
  - NVIDIA GPU CUDA compatible (optional but faster prediction)

ACORBA supports the use of GPUs for deep machine learning predictions within the TensorFlow ecosystem. Running ACORBA without a compatible GPU or no GPU (CPU integrated graphics) will produce this message in the console and the prediction will run on your CPU:

```
2021-05-04 12:25:10.634280: W tensorflow/stream_executor/platform/default/dso_loader.cc:60] Could not load dynamic library 'cudart64_110.dll'; dlerror: cudart64_110.dll not found
2021-05-04 12:25:12.018483: W tensorflow/stream_executor/platform/default/dso_loader.cc:60] Could not load dynamic library 'cudart64_110.dll'; dlerror: cudart64_110.dll not found
2021-05-04 12:25:12.019302: W tensorflow/stream_executor/platform/default/dso_loader.cc:60] Could not load dynamic library 'cublas64_11.dll'; dlerror: cublas64_11.dll not found
2021-05-04 12:25:12.020117: W tensorflow/stream_executor/platform/default/dso_loader.cc:60] Could not load dynamic library 'cublasLt64_11.dll'; dlerror: cublasLt64_11.dll not found
2021-05-04 12:25:12.031370: W tensorflow/stream_executor/platform/default/dso_loader.cc:60] Could not load dynamic library 'cusparse64_11.dll'; dlerror: cusparse64_11.dll not found
2021-05-04 12:25:12.032037: W tensorflow/stream_executor/platform/default/dso_loader.cc:60] Could not load dynamic library 'cudnn64_8.dll'; dlerror: cudnn64_8.dll not found
2021-05-04 12:25:12.032048: W tensorflow/core/common_runtime/gpu/gpu_device.cc:1757] Cannot dlopen some GPU libraries. Please make sure the missing libraries mentioned above are installed properly if you would like to use GPU. Follow the guide at https://www.tensorflow.org/install/gpu for how to download and setup the required libraries for your platform.
Skipping registering GPU devices...
```

https://developer.nvidia.com/cuda-gpus (list of compatible GPUs)

https://www.tensorflow.org/install/gpu?hl=eng (installation of CUDA for TensorFlow with NVIDIA GPUs)

#### b. Installation

Download the ACORBA.exe setup file at https://sourceforge.net/projects/acorba/ and follow the installation instructions. Check for updates and patches regularly. Your antivirus might pick up the software as a false positive (AVG antivirus does). An easy way to disable false positives (which could interfere with your analysis) is to declare the installation folder as an exception in your antivirus.
The following .exe file are part of the software and should be declared as false positive if detected by antivirus:

ACORBA.exe
testsegmentation_micro{version}.exe
testsegmentation_scanner{version}..exe
scanner{version}.exe
microscope{version}..exe
utils.exe

### 3. Image pre-processing

Don't modify the pictures either in the scanner software during scan (denoising, sharpening….) or in ImageJ except for cropping and removing of bubble etc. The models were trained on raw image so the accuracy goes down with modification as the interface root/background is different than the one the model were trained on.

#### a. Microscopy pictures

***QUICK VERSION***

***One root only in brightfield, 16-bits grayscale, >256 pixels on one side, >1 slice***

- Microscopy images must be brightfield pictures obtained by either a confocal or a widefield microscope.

- The models were trained to not recognize root hairs and detached root caps

- Image must be 16-bits scale of gray. If your images are 8-bits, the deep machine learning prediction will work but the traditional method will not.

- Images must be stacks of individual root bending/growing over time (minimum 2 timeframes) save as .tif files. It is not recommended but if you have only one time frame, duplicate it in ImageJ to have a two timeframes stack and analyze it without normalization.

- If two roots are visible in one or several frame, please save them as two different files or crop out the root of interest.

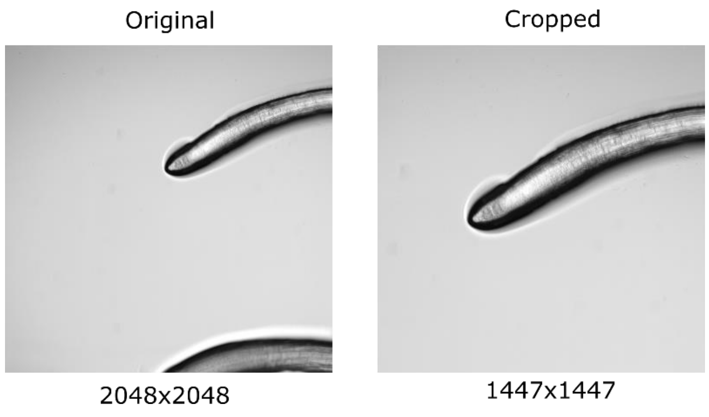

- Cropped and non-squared images will be padded (filled with mean gray value to a squared image) automatically during the analysis.

- The minimum input size must be at least 256 pixels on one side. This size corresponds to the input of the prediction models and the traditional image segmentation.

#### b. Scanner pictures

***QUICK VERSION***
***One row of seedlings only without cotyledons/hypocotyls, ROOT TIPS ON THE RIGHT SIDE, 8-bits grayscale, >256 pixels on one side, >1 slice***

- Scanner pictures must be gray pictures of individual rows of seedlings with a minimum resolution of 1200 dpi. Not respecting this resolution (higher or lower) will lead to loss of precision in the angle measurements or inaccurate prediction by the machine learning model.

- The model was trained to not recognize hypocotyls, and cotyledons but for guarantying the accuracy please remove them as described below. Bubbles, scratches, and various smears including marker pen are most of the time not detected as roots.

- Images must be stacks of one row of seedling bending/growing over time (minimum 2 timeframes) save as .tif files. It is not recommended but if you have only one time frame, duplicate it in ImageJ to have a two timeframes stack and analyze it without normalization.

- Images must be pre-processed as indicated below:
  Procedure with ImageJ or Fiji after importing all your timeframe and transformed them as a stack (Image>Stacks>Images to stack).

- If two roots are touching, please cropped them out of the images as the software was not programmed to deal with touching elements.

- The minimum input size must be at least 256 pixels on one side. This size corresponds to the input of the prediction model.

Original stack

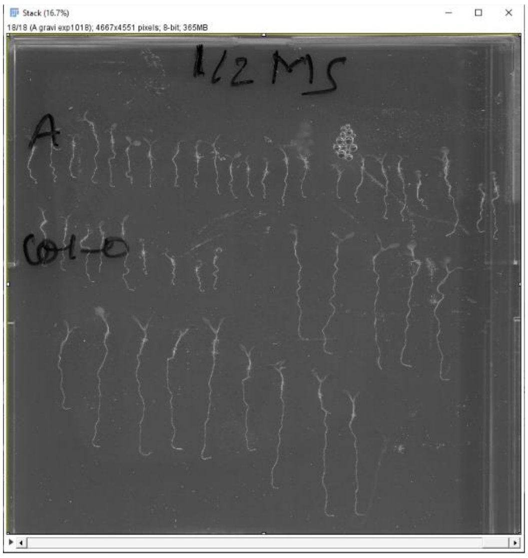

Flipped and turn 90° stack
(root tips must be on the right)

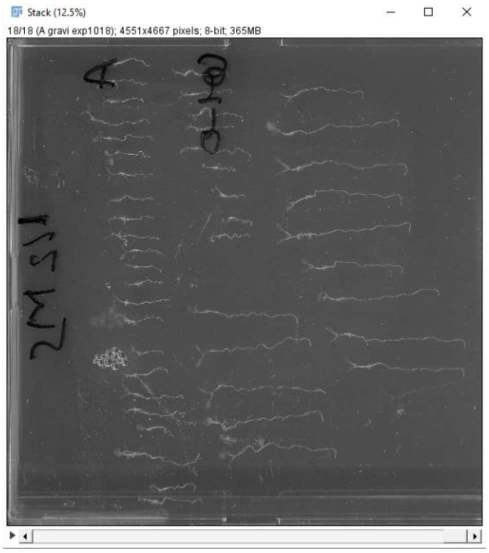

Duplication of a single row
rectangular selection and
Ctrl+Shift+D

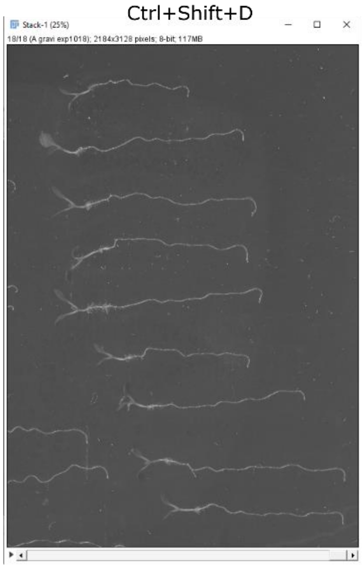

Polygonal selection to remove cotyledons
and bubble, big scratchs...

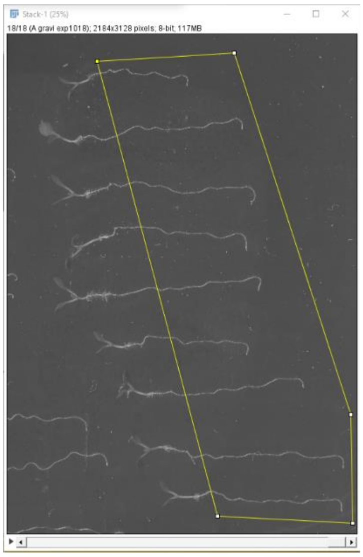

Edit>Clear Outside
with black pixels

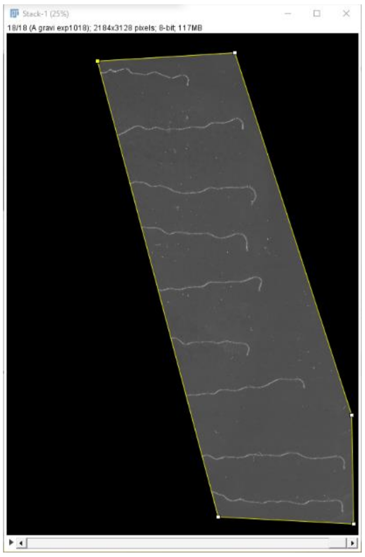

Second crop to reduce file size
and processing time. Save as .tif

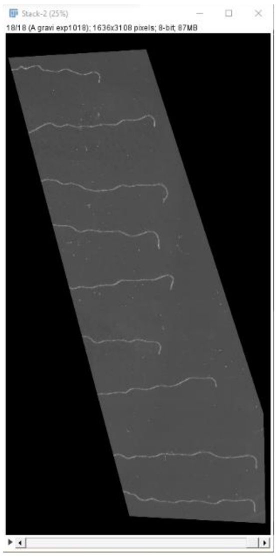

### 4. Quick start

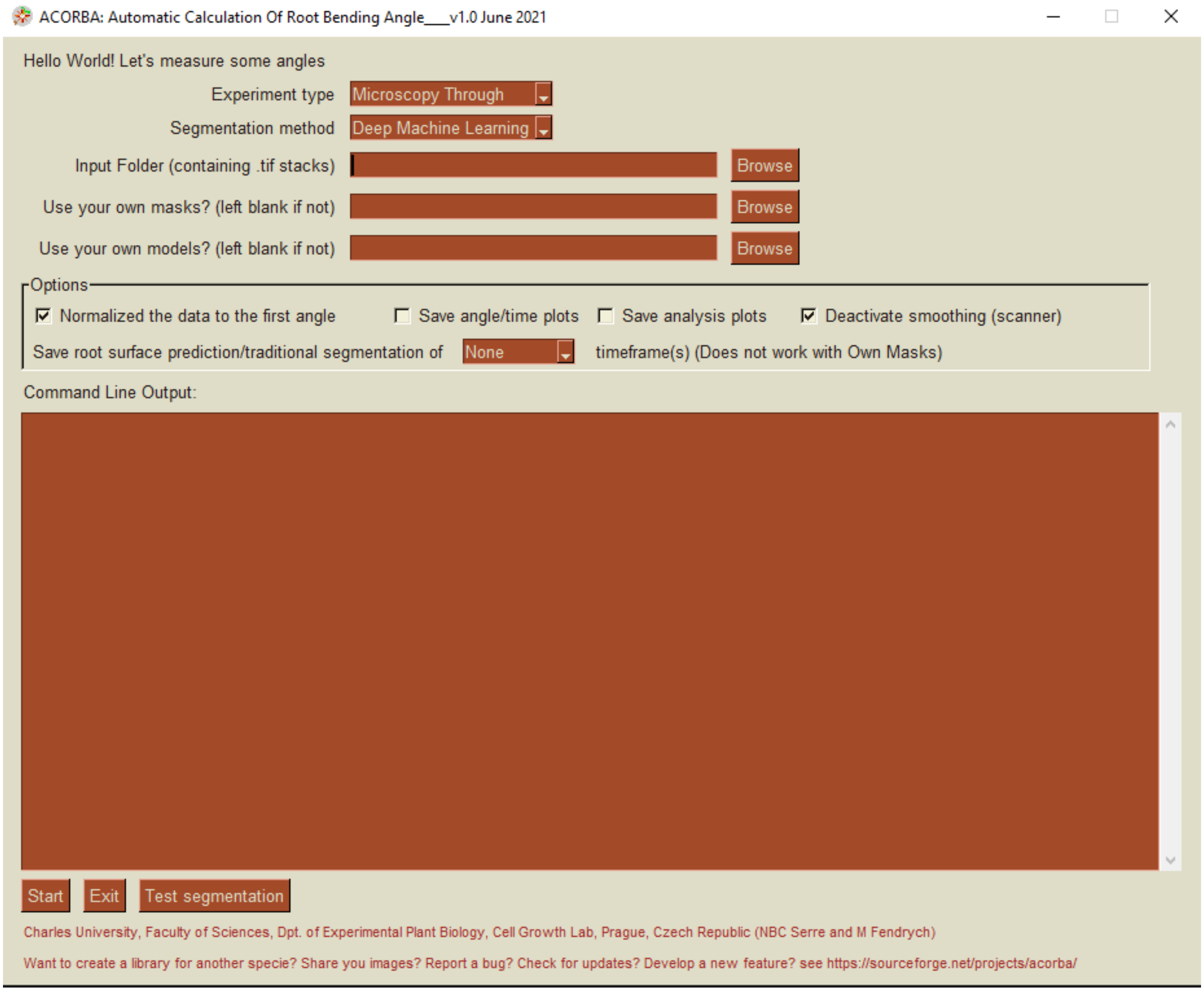

ACORBA Graphic User interface

### 5. Modes and options

#### a. Experiment type

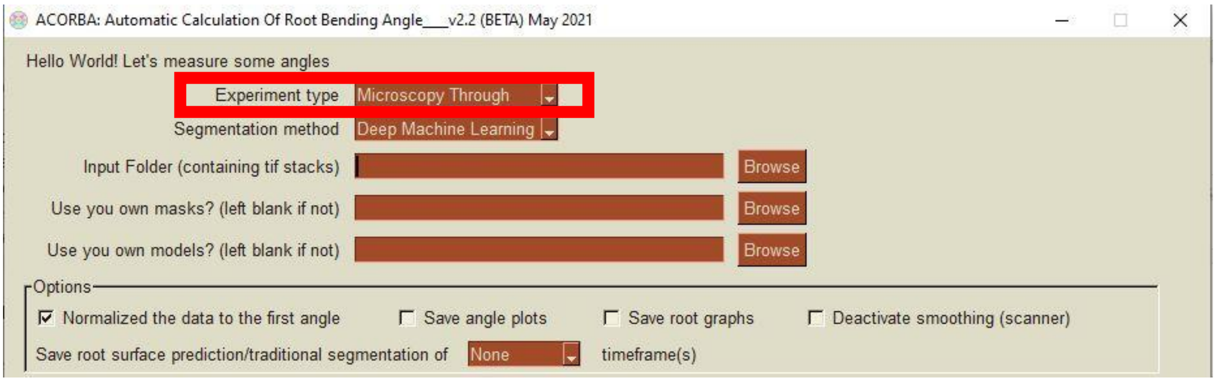

- Microscopy Through

This experiment type corresponds to roots bending on a vertical stage microscope with the root on top of a thin layer of ½ MS medium. In these conditions, the root is blurred but is less mechanically stressed than with the sandwich method.

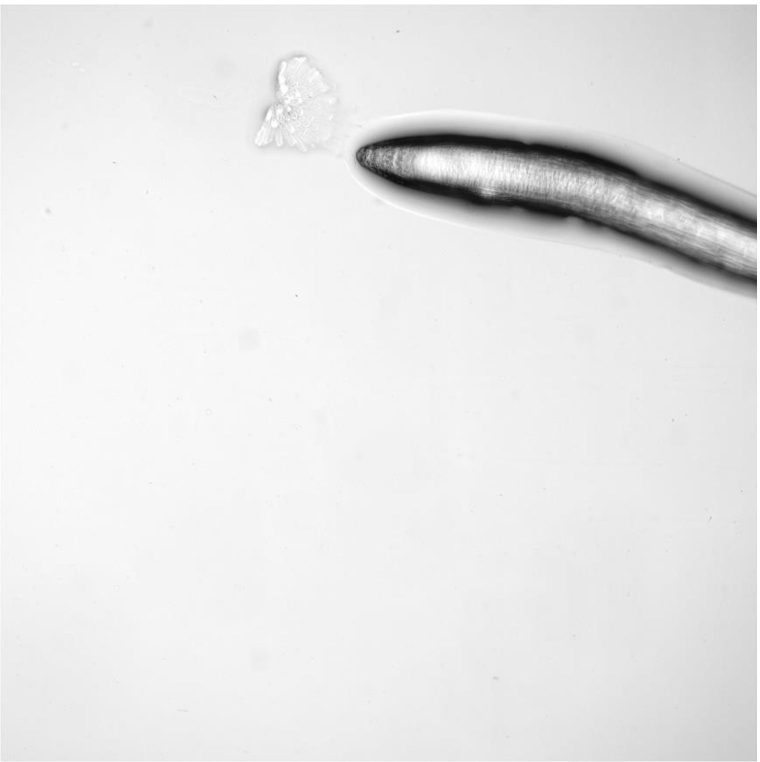

- Microscopy Sandwich

This experiment type corresponds to roots bending on a vertical stage microscope with the root sandwiched between the chamber coverglass and a layer of ½ MS medium.

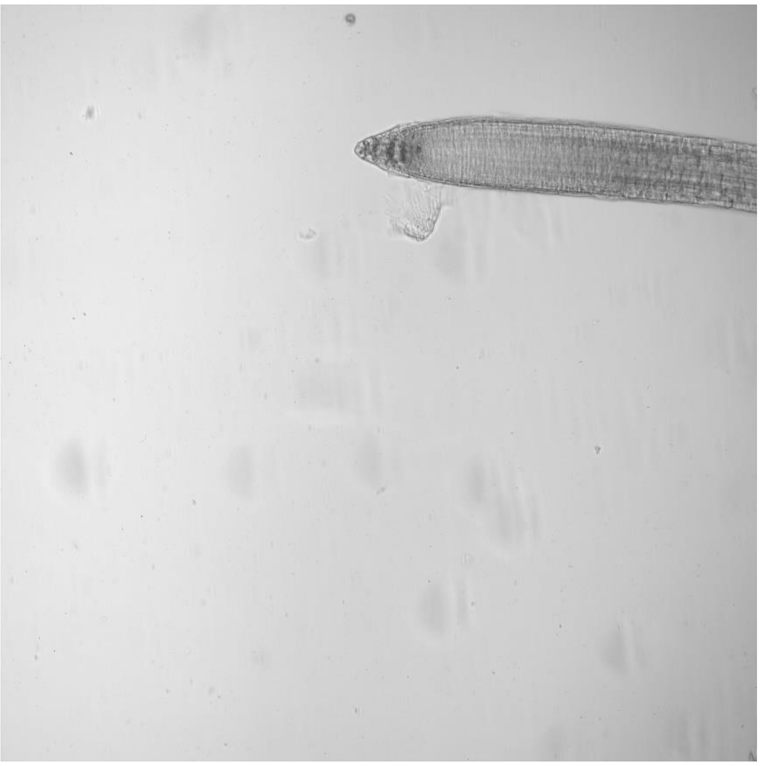

- Scanner

This experiment type corresponds to plates scanned on a flatbed scanner over time.

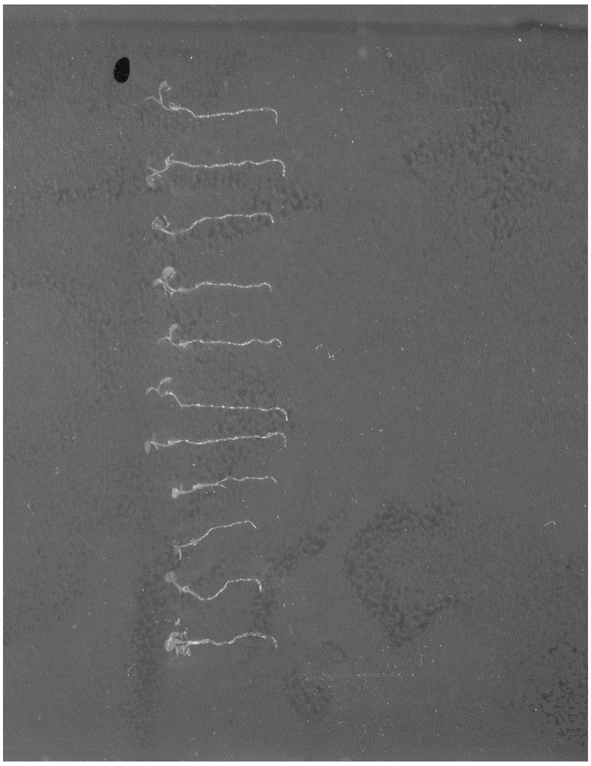

#### b. Segmentation method

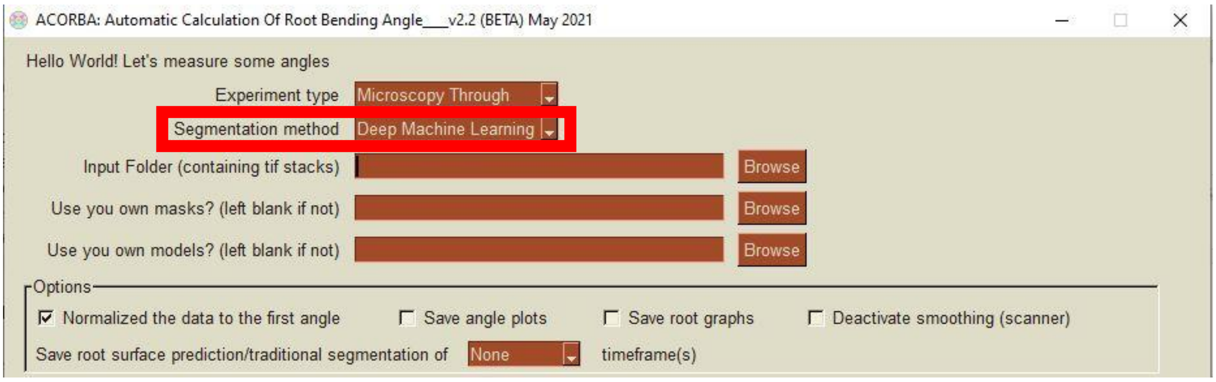

- Deep Machine Learning

This method of segmentation uses deep machine learning image segmentation. Prediction models were developed using hundreds of roots from various inputs to create models that can accurately predict root surfaces/tips. The models were trained to avoid recognizing unwanted elements such as cotyledon and hypocotyls, bubbles, scratches, root hairs and floating root cap.

- Traditional

This method of segmentation uses a traditional approach to images segmentation. See Original paper or code in the utils.py script for details

#### c. Use your own segmentation masks?

To create the masks, see paragraph 6. Once the masks are prepared as requested, precise where is the folder containing them in the software.

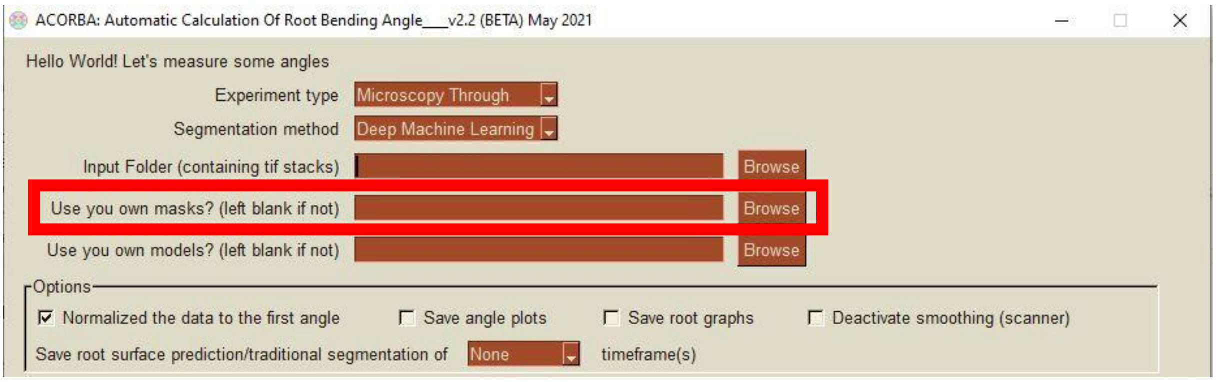

#### d. Use your own models and weights?

See paragraph 7. Once the models and weights are prepared as requested, precise the folder containing your models and weights in the software.

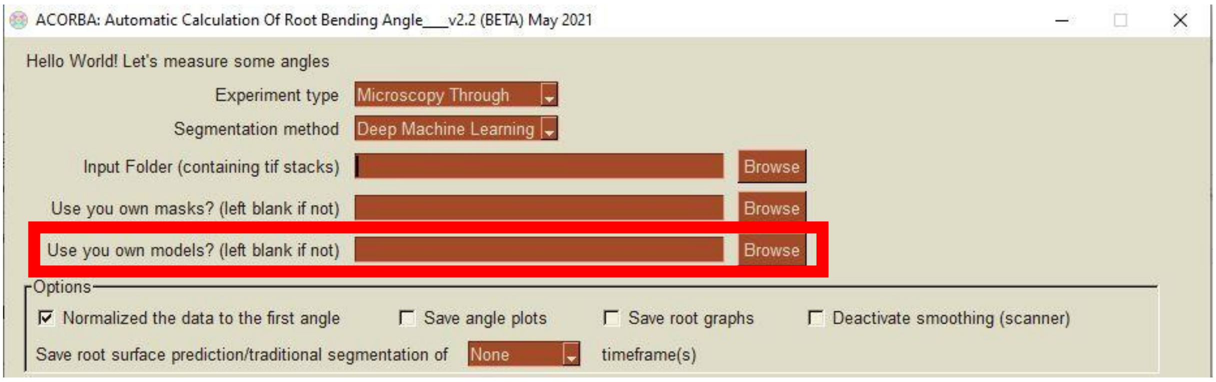

#### e. Normalized data to the first angle

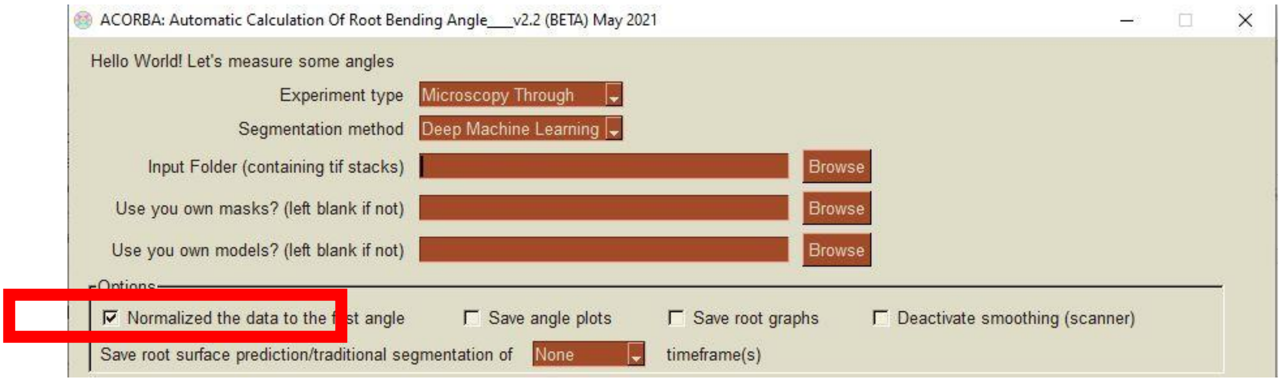

Normalization of the data the first angle is to calculate the relative angles over time. This option is ticked by default and will subtract the first angle of a root to the following angles. For example:

| Timeframes | Raw data | Normalized |
|---|---|---|
| 1 | 5° | 5-5=0° |
| 2 | 10° | 10-5=5° |
| 3 | 15° | 15-5=10° |
| 4 | 20° | 20-5=15° |

When this option not ticked, the lists of angles will be exported as an excel file with only the raw measurements in one sheet.
When this option is ticked, the lists of angles will be exported as an excel file with two sheets: Raw data and Normalized data.

#### f. Save angle/time plots

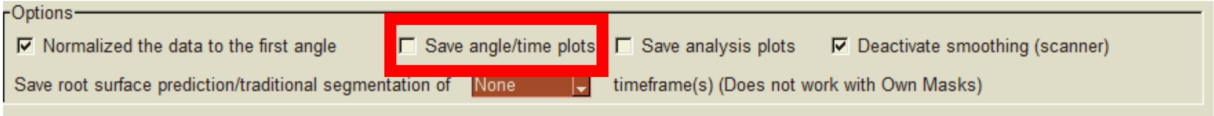

When this option is ticked (unticked by default), the software will automatically save the angle over time plot of one root (microscopy) or a row of roots (scanner). If the normalized option is ticked, the normalized plot will be saved.

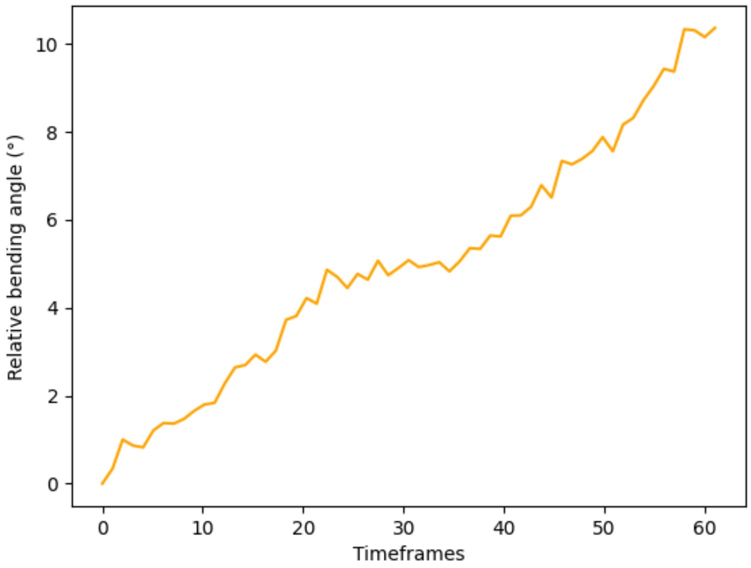

#### g. Save analysis plots

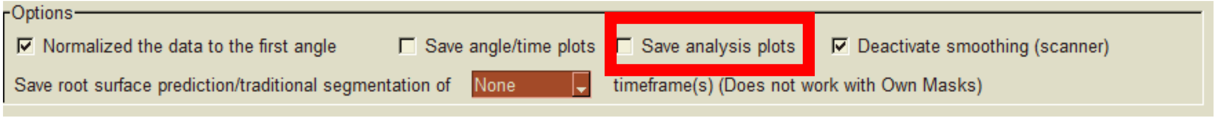

When this option is ticked (unticked by default), the software will automatically save the plots that are produced during the analysis for each root/row of roots and each timeframe. The pictures are named after your original file name and contains the name of the file, the timeframe number and the angle calculated.

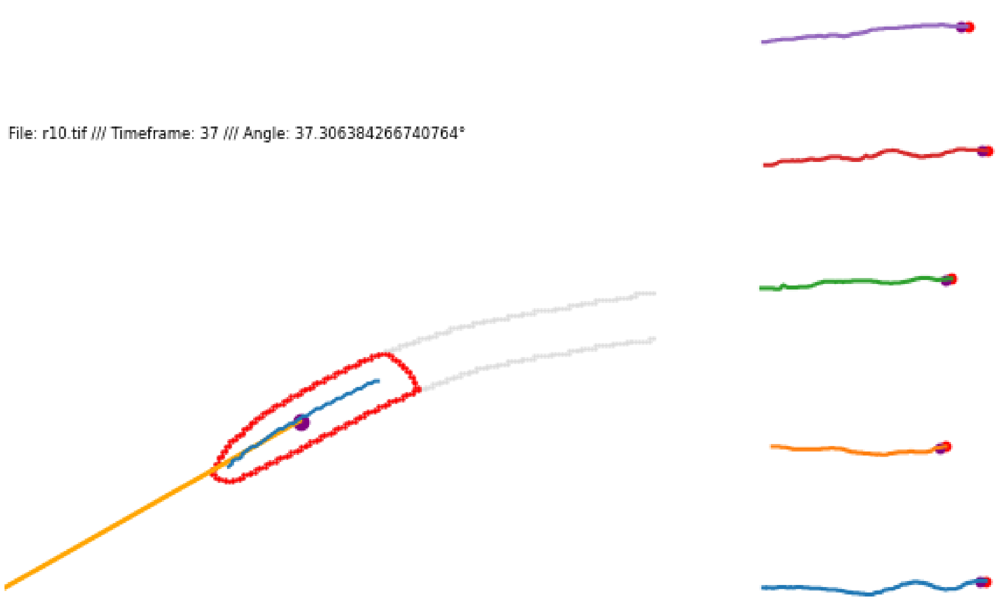

#### h. Save root surface predictions

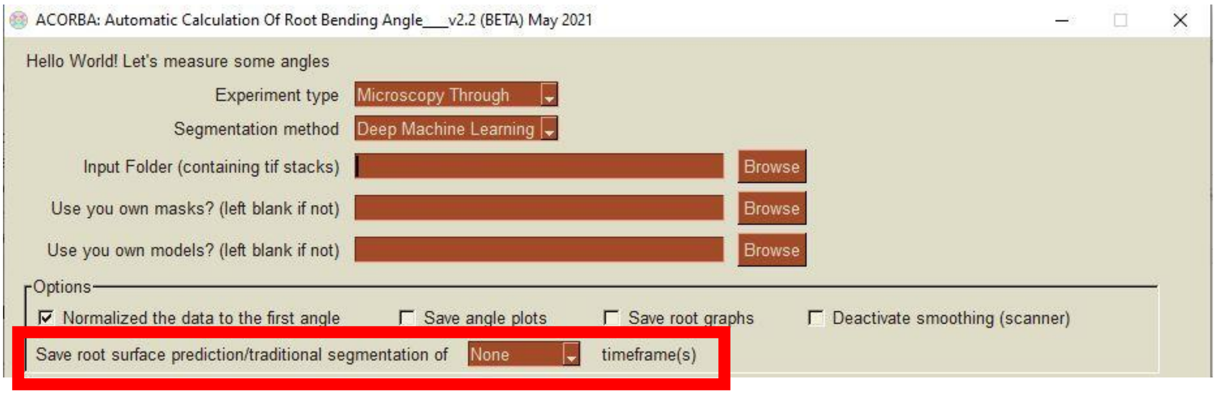

When ticked the software will automatically save the predictions done for your roots for all the timeframes. For microscopy you will have two files corresponding to the root tip and root surface predictions (see next page). In every case there will be 3 columns corresponding to the original picture, the segmented picture, and the overlay between the two first.

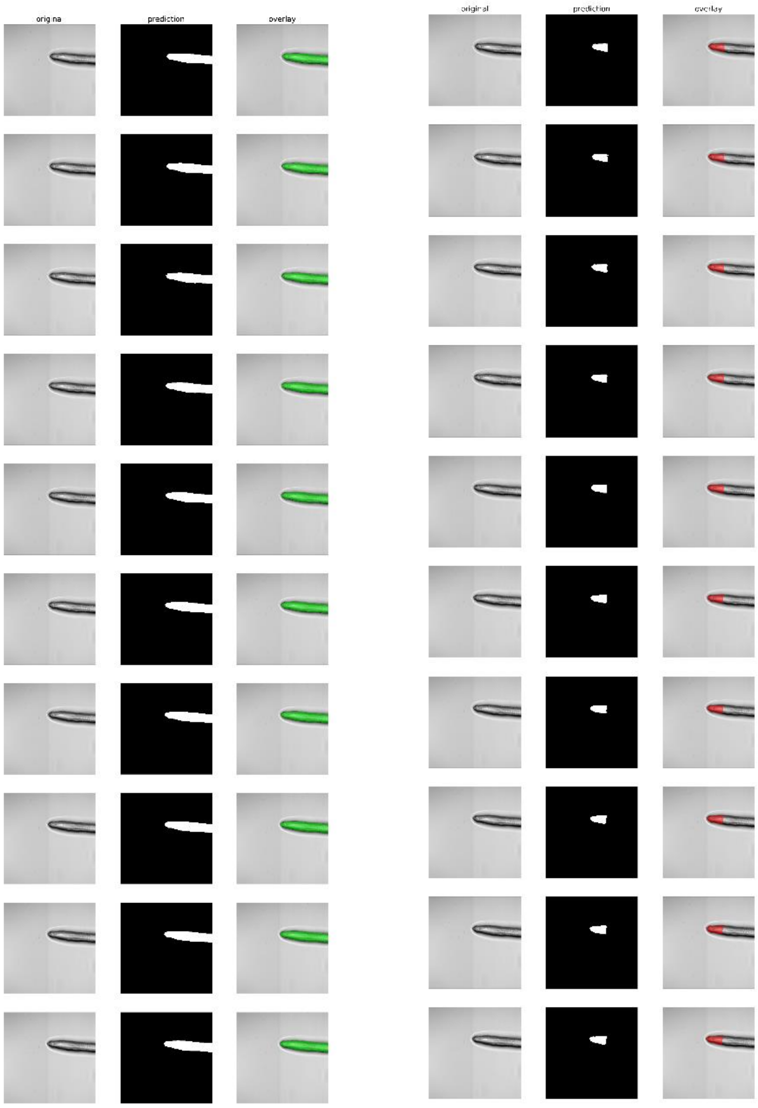

#### i. Deactivate smoothing (scanner)

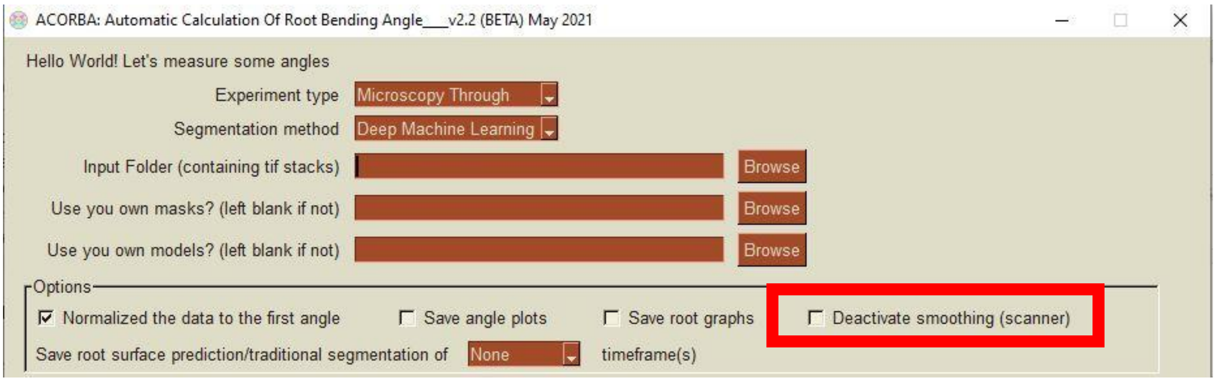

Smoothing by a rolling average factor 2 can be applied (**not ticked by default**) to scanner data (even the raw data exported). You can enable this parameter by ticking the box.

#### j. Test segmentation

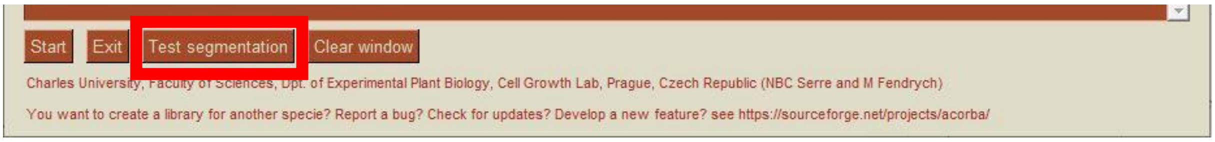

Test segmentation should be the first analysis you carry out when you want to analyze a new dataset. It exports the comparison between the machine learning and traditional methods of segmentation into a multi tiles picture (automatically saved in your folder). This test allows you to determine which method is the best for your dataset and in some cases to remove roots or modify them before the actual angle analysis.

#### k. Start

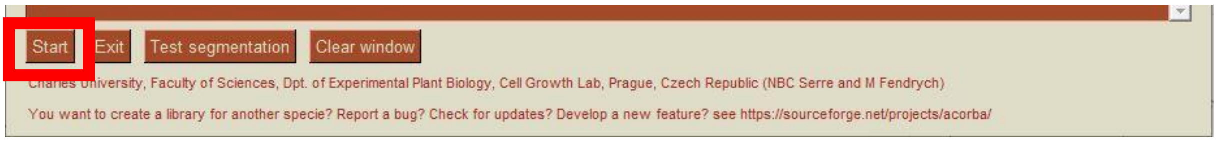

The start button is the button to start the analysis of all the files present in the specified folder. The analysis will be carried out using the parameters specified above.
At the beginning of the analysis the console will recapitulate the parameters provided.

For example:

*Root folder: D: /Experiments /*
*Save angle plot: False*
*Save root plot: False*
*Experiment type: Microscopy Through*
*Normalization: True*
*Save prediction plot: None*
*Segmentation method: Deep Machine Learning*
*Use masks from:*
*Use models from:*
*Deactivate smoothing: True*

#### 6. Own segmentations masks

##### a. Recommended methods to annotate roots

- ImageJ
- Apeer.com, Zeiss free online image analysis platform

##### b. Required format and way to proceed

Segmentation masks must be binary tif stacks with a background value of 0 and the root surface annotated with a value of 255 or 1.

**Quick tips to segment Scanner pictures with ImageJ FIJI**

Use subtract background (with, for example, the settings below) and other image processing to enhance the thresholding

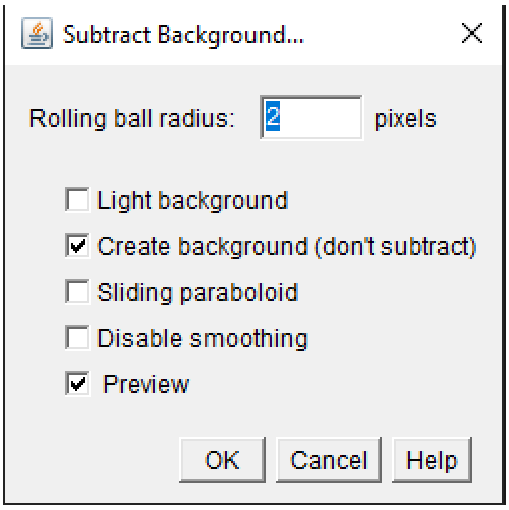

Threshold with default or Huang method are showing accurate results. Be sure that your roots are not segmented in several pieces. If some are, re-threshold or draw the space missing with the brush.

**Quick tips to prepare surface and tip binary pictures with ImageJ FIJI**

- Roots in microscopy are usually not hard to threshold but sometimes the exposure can change a little and make the thresholding of the whole stack a nightmare. Use a simple bleach correction with default settings to remediate this.

- If you have root imaged with our Microscopy through method, threshold the outer root and then fill the middle manually.

- Once you have the surface segmented, save it with the same name as the original in a folder named “surface”.

- To prepare the root tip masks efficiently, reopen your surface segmentation and Image>Stack>Stack to Images. Then, roughly select the root tip with a rectangular or polygonal selection (see image below) and Edit>Clear Outside (be sure your color are set as shown below). You can set Clear Outside as a keyboard shortcut in ImageJ to make the process faster. Once all the root tips are done, re-stack yours images (Image>Stack>Images to Stack) and save it in a folder intitled “tip”.

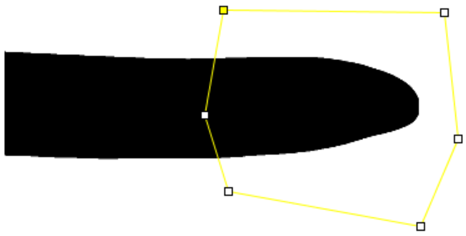

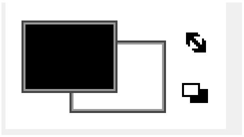

The masks must have the same names has the original pictures and be placed into a separated folder. For Microscopy, they need to be separated in two folders but still with the same name as the original stack:

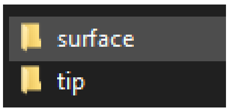

Indicate “Own masks” has a segmentation method and specify the folder path in the software under “Use your own masks”.

### 7. Custom models

#### a. Create a new model with us?

- General requirements

If you wish to develop a Deep Machine Learning models and weights in collaboration with us, please open a ticket at https://sourceforge.net/projects/acorba/ .
Note that to create a model from scratch you will need a bare minimum of 100 annotated roots images. For microscopy both root tip and root surface need to be annotated.

#### b. Overview of the way to proceed

- Models and weights with TensorFlow

ACORBA machine learning predictions are implemented with the modules TensorFlow and Keras. Follow the steps below to implement them in ACORBA.
Once you have established your library of images and masks, please adapt the python script “Image_preprocessing_microscopy.py” or “Image_preprocessing_scanner.py”.
Then, you can train your model of choice and save your model and weights to .json .h5 and, respectively:

```
from keras.models import model_from_json
# serialize model to json
json_model = model.to_json()
#save the model architecture to JSON file
with open(MyModel.json', 'w') as json_file:
        json_file.write(json_model)
#saving the weights of the model
model.save_weights('MyModel_Weights.h5')
```

Be aware that if you want to use ACORBA has it was originally programmed, your libraries for microscopy pictures need to be 16 bits and padded/resized to 256x256 pixels and scanner pictures need to be 8 bits and padded to a size dividable by 256 tiles then divided into 256 tiles. Please see and adapt the python script “Image_preprocessing_microscopy.py” or “Image_preprocessing_scanner.py”.

If you are a Python enthusiast, feel free to modify the original code to your needs. The Jupyter notebooks used in Google Collaboratory to train our models are available on the Sourceforge repository. If some code is not clear, open a ticket on the Sourceforge. According to the license, if you use entire or partial parts of this code, please cite the original ACORBA paper.

Note that for the microscopy method you will need both a model and weights to predict root tip and root surface. Models and weights names need to contain “tip” and “surface” (example below) (even if the model for tip and surface is actually the same, create a copy). For scanner, the names are not important.

model_surface_through_256_sat_jaccard_update16062021_54.h5
model_through_tip_256_sat_jaccard_update17062021_176.h5
surface256.json
tip256.json

Call them in ACORBA (see 5.d) by indicating the folder where they are stored.

- Model from the online platform Zeiss Apeer (apeer.com)

Unfortunately, it is not possible to reuse model created by the online platform Apeer as they use custom layers in the model creation. If you have a working model from this platform, you could still use their interface to segment your images and use the own segmentations method in ACORBA (see paragraph 6.)

### 8. Measure root waving

Root waving can be measured from scanned plates. The images must be pre-processed has described in paragraph 3.b. Put the root tips on the right side of the picture.
Positive values correspond to angle going left and negative value to angles going right.
Be aware that ACORBA only provides the waving angles and not the length, VGI or HGI parameters. These parameters were not implemented has it was not the main goal of the software but they are extractable from the images. If you are interested in these parameters, please contact us by opening a ticket or a forum thread at https://sourceforge.net/projects/acorba/ .

### 9. FAQ and Troubleshooting

**Before opening a forum thread or ticket on the repository please read this FAQ.**

**Be aware that the maintenance of this software is only done by one person and that it's not my main research activity.**

#### a. Missing DLLs

Please install latest Microsoft Visual C++ redistributable x64, if still not working take a screenshot of ACORBA's console and open a thread or ticket at https://sourceforge.net/projects/acorba/

#### b. Missing rolling_ball.cy

Please install Microsoft Visual C++ redistributable x64.

#### c. Access denied for .xls output file to be written

Try running ACORBA as an administrator or you may not have the rights to write on the drive containing the folder (e.g., folder on a shared server with reading rights only).

#### d. "Index out of bound"

```
Oups something went wrong!
index 0 is out of bounds for axis 0 with size 0
```

This error happens most of the time when your roots (or just one is too short in scanner)> Cropped them longer. It can also happen when big bubbles on plates are detected > Removed them in pre-processing.

#### e. Deep machine learning is not good, traditional also

Be sure that you have not modified the raw pictures in any way except cropping. The models were trained on raw pictures and thus, any modification would be confusing for the predictions.

Microscopy pictures needs to be 16 bits and scanner 8 bits.

Sometimes just flipping the root stack on the horizontal axis can enhance (or decrease) accuracy.

If none of these solutions work, you can refer to paragraph .6 to prepare your own binary masks.

##### f. I have two root tips predicted on one root

If the second root tips (mis-predicted one) is smaller, the actual root tip will be selected not the second one. Otherwise try to modify slightly the root where the second one is predicted in order to still have a correct surface prediction but not a second root tip. A little square with the color of the background can work.

##### g. Extremely long time to start the angle calculation

This can happen for several reasons including loading pictures from a remote server or an external drive. It is recommended that the folder you want to analyze is present on the same drive as the ACORBA installation, you will also have better loading times if your drive is an SSD drive.

This can also happen if your CPU does not have enough computing power. In this case the calculation of angles itself will also be slow.

##### h. [0x7FF952949FE0] ANOMALY: meaningless REX prefix used

This error is not debugged for the moment. It very rarely appears when you finish an analysis, but it does not influence the calculation or the output export.

##### i. WARNING:tensorflow:AutoGraph could not transform <function Model.make_predict_function.<locals>.predict_function at 0x000001D46681B700> and will run it as-is.

```
WARNING:tensorflow:AutoGraph could not transform <function Model.make_predict_function.<locals>.predict_function at 0x000001D46681B700> and will run it as-is.
Please report this to the TensorFlow team. When filing the bug, set the verbosity to 10 (on Linux, `export AUTOGRAPH_VERBOSITY=10`) and attach the full output.
Cause: Unable to locate the source code of <function Model.make_predict_function.<locals>.predict_function at 0x000001D46681B700>. Note that functions defined in certain environments, like the interactive Python shell do not expose their source code. If that is the case, you should to define them in a .py source file. If you are certain the code is graph-compatible, wrap the call using @tf.autograph.do_not_convert. Original error: could not get source code
To silence this warning, decorate the function with @tf.autograph.experimental.do_not_convert
```

This error pops up in the console during the loading of the TensorFlow library. No fix has been found so far but it does not influence the analysis.

##### j. My segmentation is getting less accurate over time

This problem can be experience when the background/exposure of the root is changing overtime. Sometimes a simple bleach correction with default parameters in ImageJ is solving the problem.

#### k. Other

If you are running into any other problem, please open a ticket at https://sourceforge.net/projects/acorba/. We will get back to you as soon as possible. Be aware that only one person maintains this software so answering time may vary.

#### l. The graph window keeps bothering me while I do something

For scanner, you will need to reduce it for every plate.

For microscopy reduce it only one per analysis.

The right time to reduce this window is when it is active, meaning during the angle calculation when the graphs are actualizing.

### 10. Running/Modifying ACORBA Python code

All the:

- python scripts to run ACORBA,
- Jupyter notebooks used to train the models
- Training libraries
- Test examples

Are available at https://sourceforge.net/projects/acorba/

According to the license, if you use entire or partial parts of this code, please cite the original ACORBA paper.

The following Python modules and libraries are required to run ACORBA:

- Scikit Image
- Scikit Learn
- TensorFlow
- Keras
- Keras-segmentation
- OpenCV
- PySimpleGUI
- Matplotlib
- Pandas
- Numpy
- CV2 rolling ball
- Fil Finder
- Astropy

### 11. Citations

This work was developed by Nelson BC Serre and Matyáš Fendrych at Charles University, Faculty of Science, Department of Experimental Plant Biology, Cell Growth Lab., Prague, Czech Republic.
The ACORBA software hosted on SourceForge is maintained by Nelson BC Serre.
According to the license, if you are using entire or partial parts of this code, please cite the original ACORBA paper.
