## Supplemental figures 2, 6, 7 for "ACORBA: Automated workflow to measure *Arabidopsis thaliana* root tip angle dynamic"

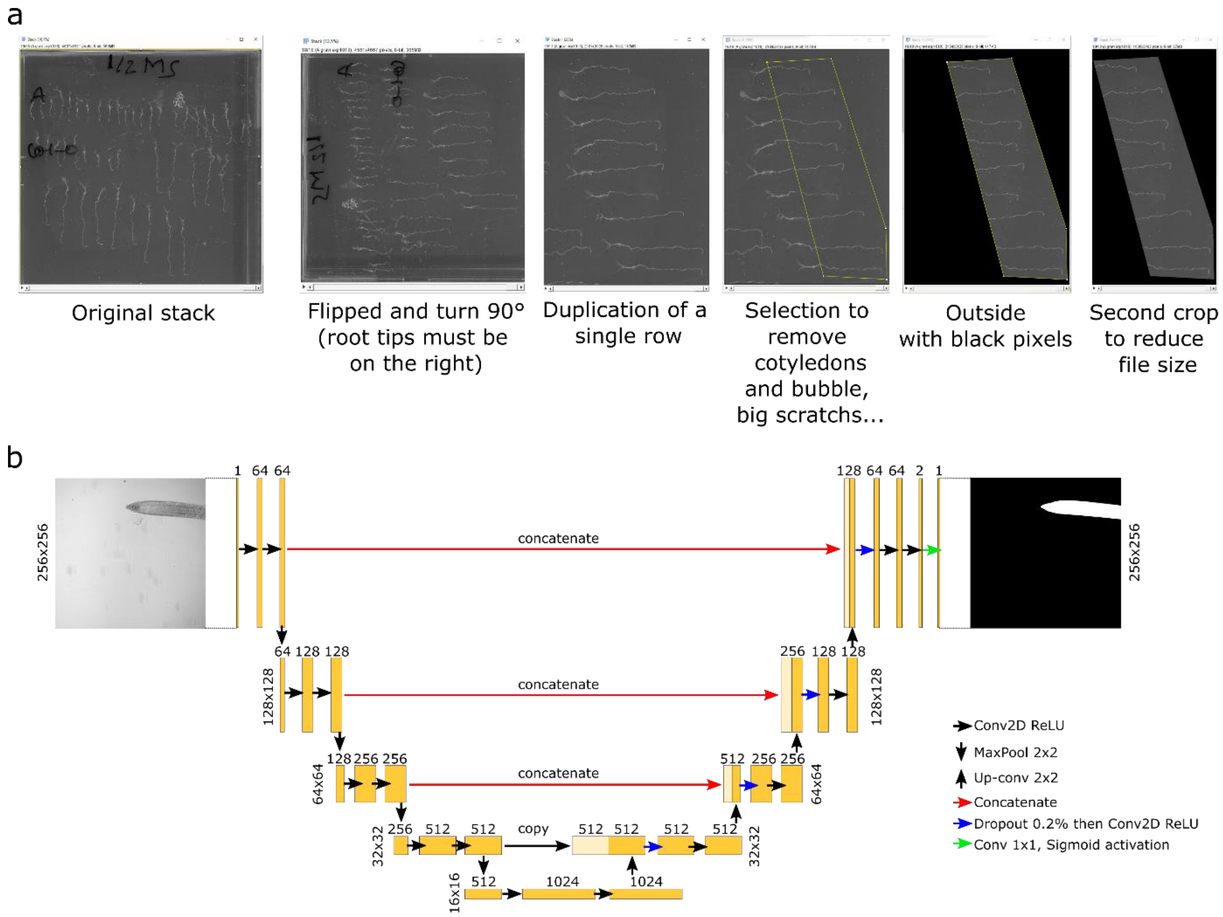


***Figure Supplemental 2:*** *(a) Image preprocessing for flatbed scanner images. First, the raw images from the scanning process are stacked into a .tif. Then, the root tips are placed on the right side of the image. Individual rows are isolated and finally cotyledons and hypocotyls are cropped out of the image. (b) UNET architecture of the deep machine learning prediction models.*

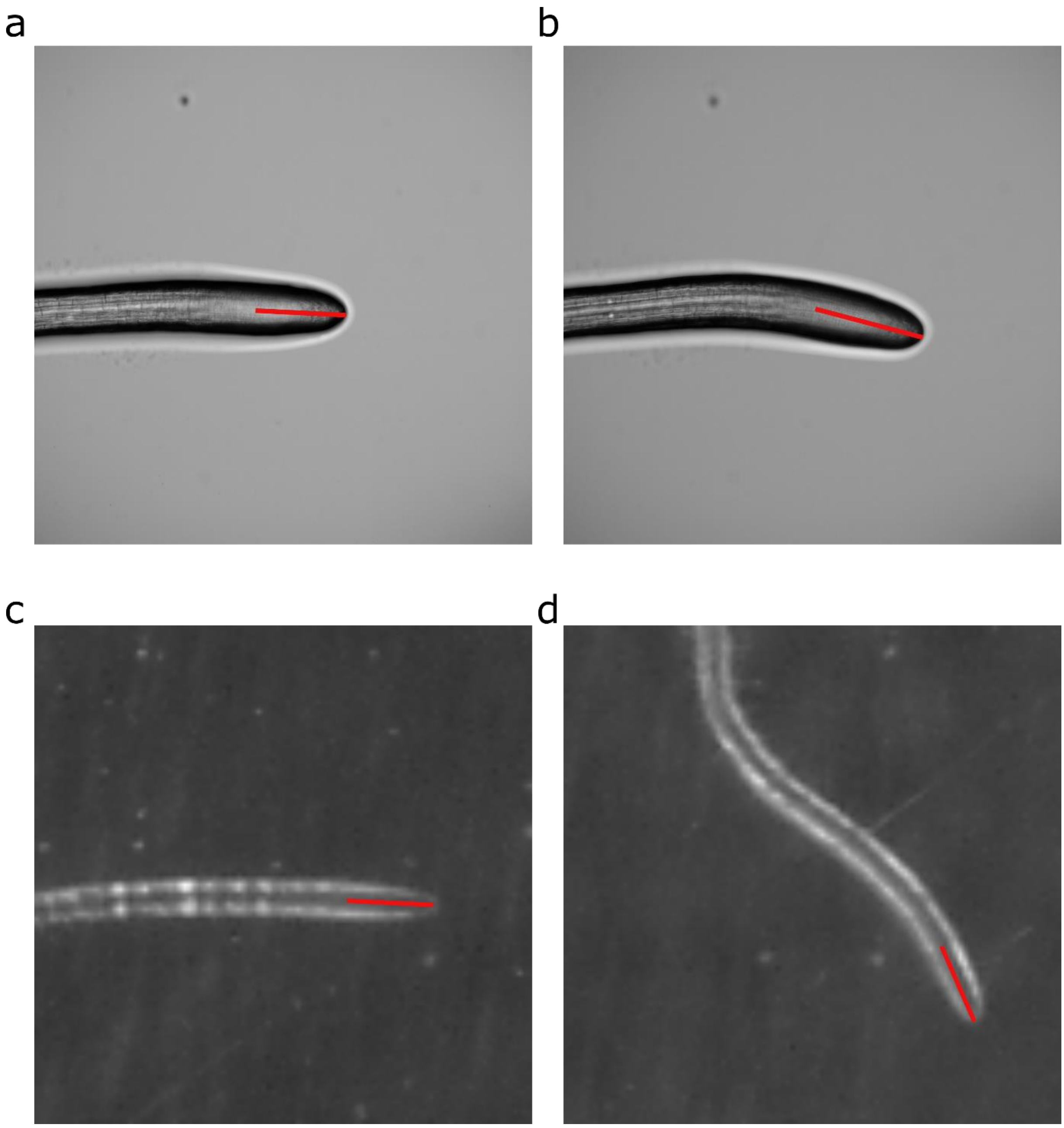


***Figure Supplemental 6:*** *Manual measurements of root tip angles using ImageJ. With the straight-line tool, a line is drawn (red line) and the angle is measured by ImageJ. (a) and (b) represent a root bending imaged with the microscopy through method in a vertical-stage microscope. (a) After 2 minutes and (b) 62 minutes of gravistimulation. (c) and (d) represent a root bending imaged with a flatbed scanner. (a) After 2 minutes and (b) 10 hours of gravistimulation.*

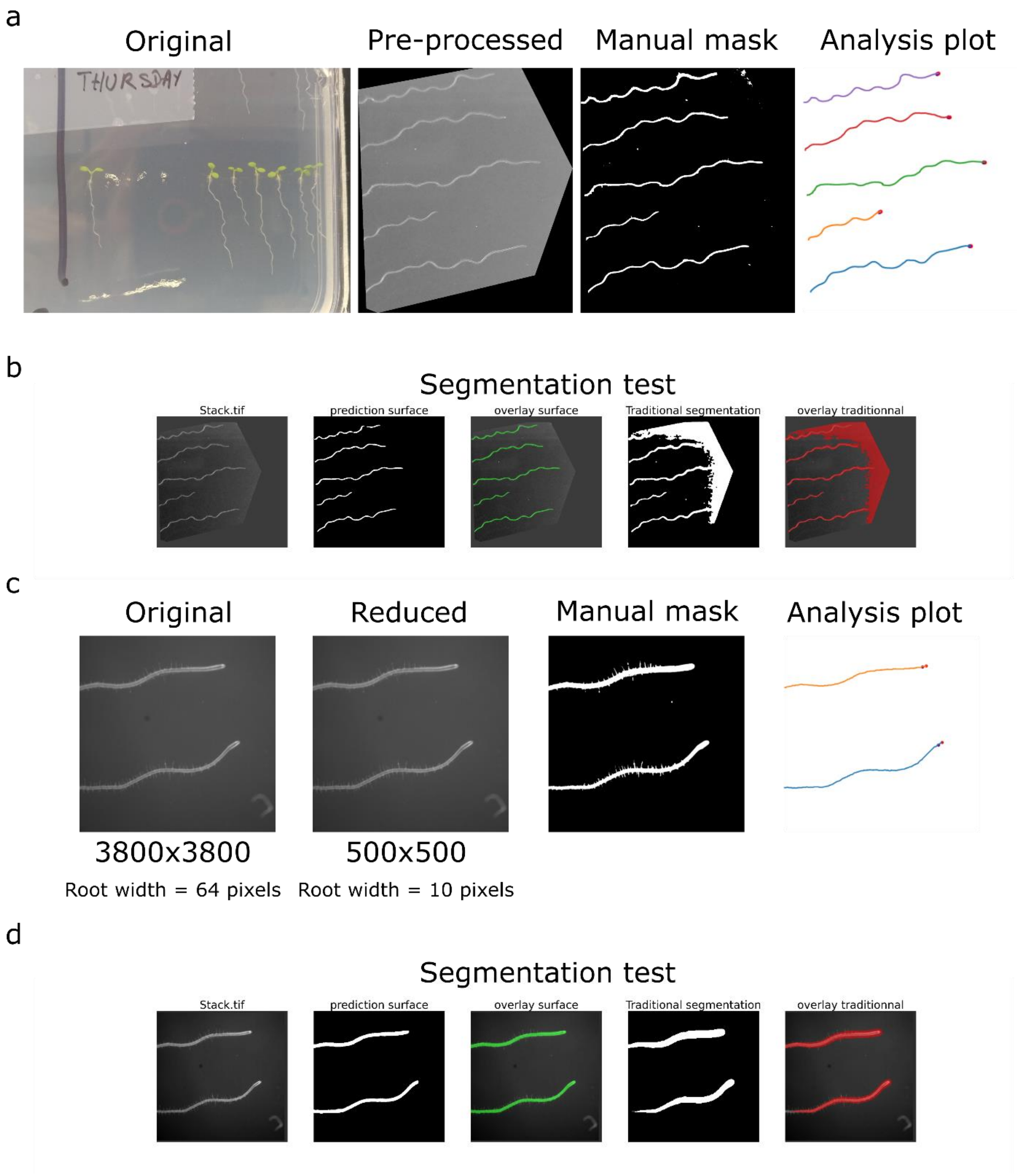


***Figure Supplemental 7:*** *(a) Semi-automated ACORBA analysis of root angle from mobile phone image. (b) Segmentation test of the mobile phone image comparing both deep machine learning and automated traditional segmentation. (c) Semi-automated ACORBA analysis of root angle from horizontally growing Col0 observed with a stereomicroscope. (d) Segmentation test of the stereomicroscope image comparing both deep machine learning and automated traditional segmentation.*
