## Supplementary figures and images for "ACORBA: Automated workflow to measure *Arabidopsis thaliana* root tip angle dynamic"

### Supplemental image 1

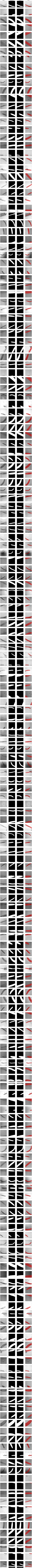

### Supplemental image 2

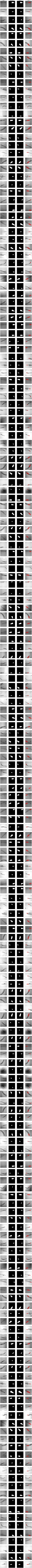

### Supplemental image 3

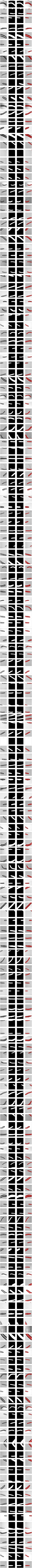

### Supplemental image 4

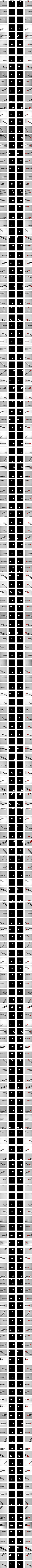

### Supplemental image 5

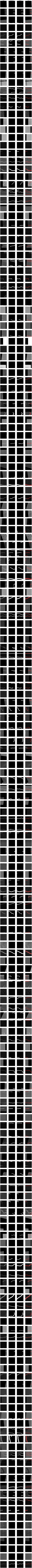

### Supplemental image 6

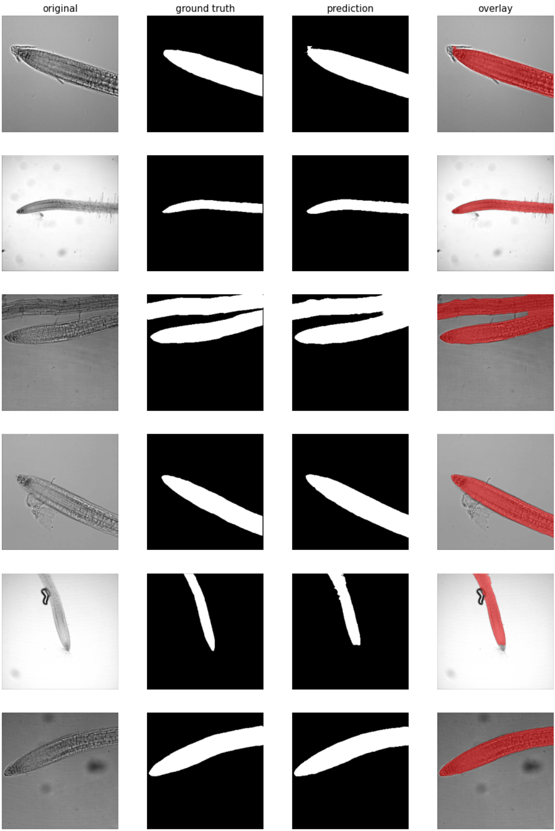

### Supplemental image 7

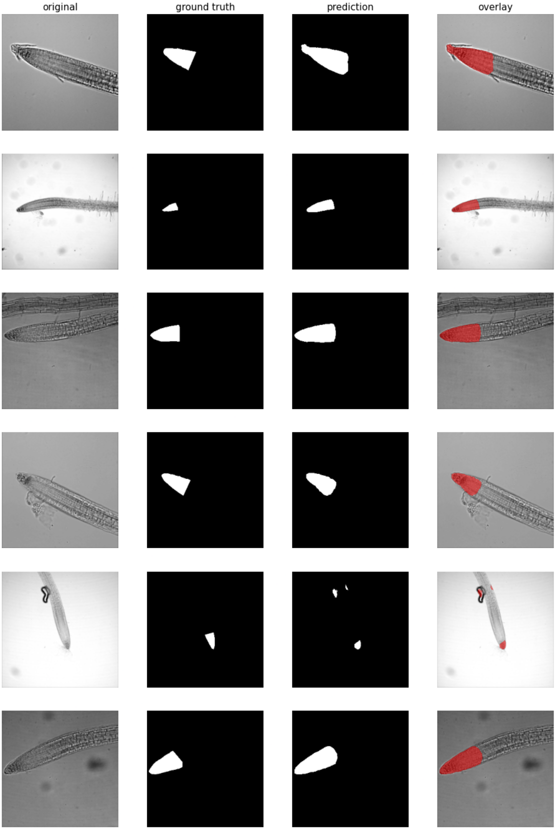

### Supplemental image 8

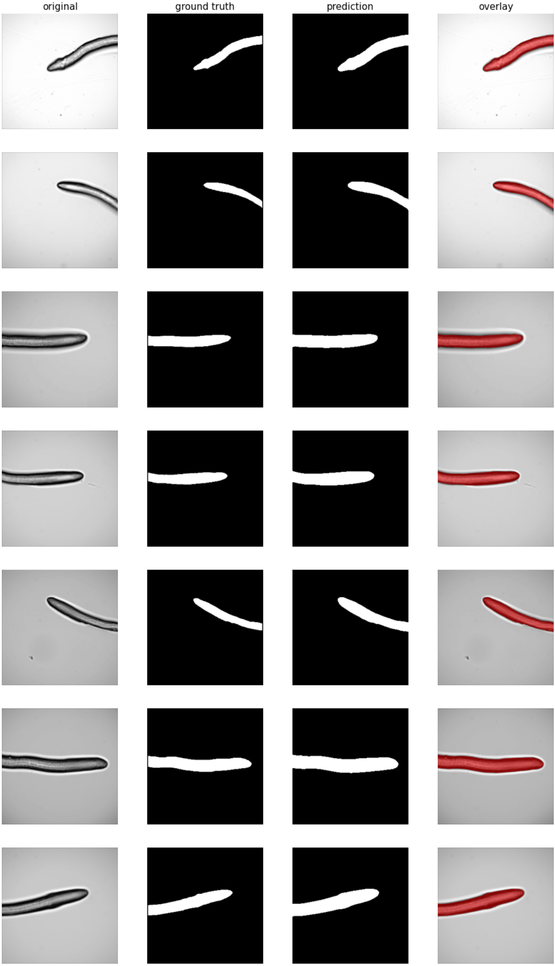

### Supplemental image 9

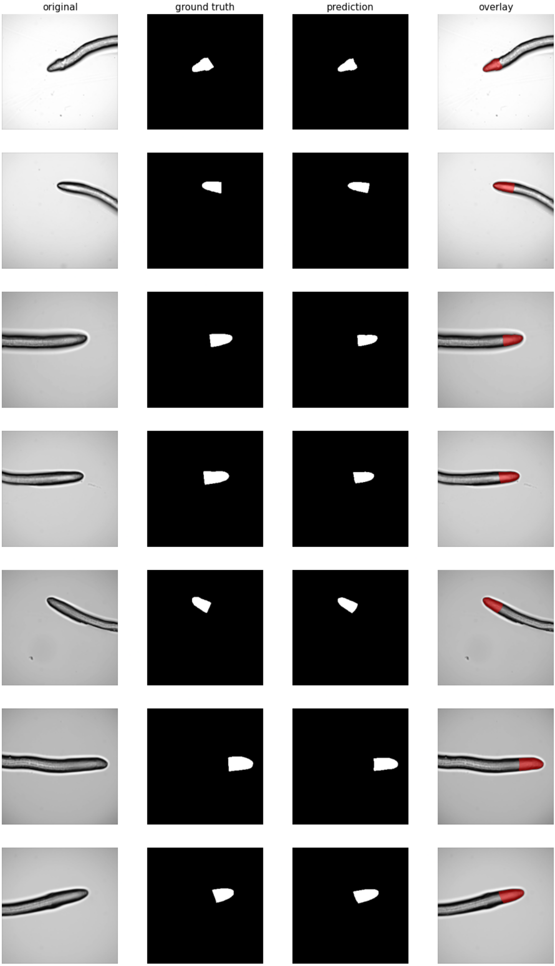

### Supplemental image 10

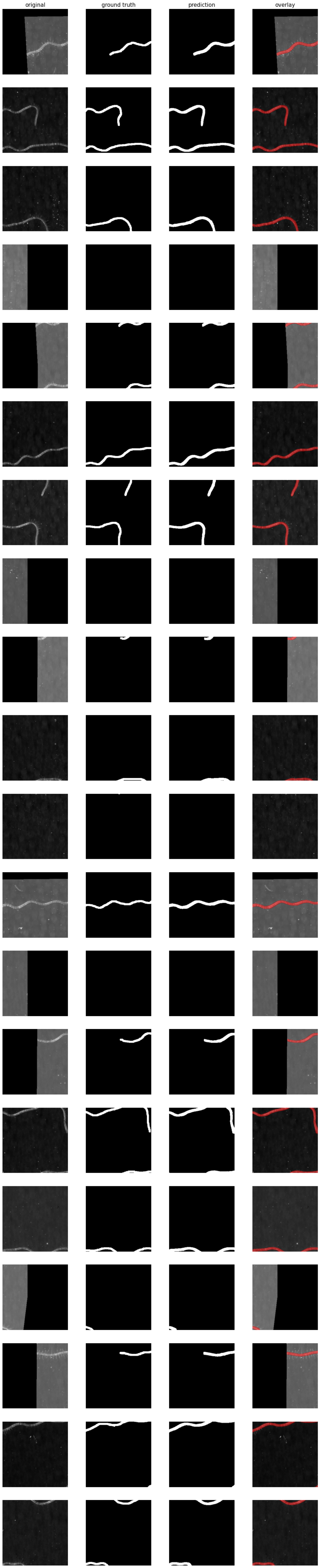
